# Dynamics of endogenous PARP1 and PARP2 during DNA damage revealed by live-cell single-molecule imaging

**DOI:** 10.1101/2022.03.12.484081

**Authors:** Jyothi Mahadevan, Asmita Jha, Johannes Rudolph, Samuel Bowerman, Domenic Narducci, Anders S Hansen, Karolin Luger

## Abstract

PARP1 contributes to genome architecture and DNA damage repair through its dynamic association with chromatin. PARP1 and PARP2 (PARP1/2) recognize damaged DNA and recruit the DNA repair machinery. Using single molecule microscopy in live cells, we monitored the movement of PARP1/2 on undamaged and damaged chromatin. We identify two classes of freely diffusing PARP1/2 and two classes of bound PARP1/2. The majority (> 60%) of PARP1/2 diffuse freely in both undamaged and damaged nuclei and in the presence of inhibitors of PARP1/2 used for cancer therapy (PARPi). Laser induced DNA damage results in a small fraction of slowly diffusing PARP1 and PARP2 to become transiently bound. Treatment of cells with PARPi in the presence of DNA damage causes subtle changes in the dynamics of bound PARP1/2, in contrast to bulk studies that suggest PARP trapping. Our results imply that next-generation PARPi could specifically target the small fraction of DNA-bound PARP1/2.

## Introduction

Repair of damaged DNA begins with the recognition of the lesion by protein factors as exemplified by the rapid detection of single and double stranded DNA breaks (SSBs and DSBs) by the nuclear enzymes PARP1 and PARP2 (Benjamin and Gill, 1980; Haince et al., 2008; Mortusewicz et al., 2007). Upon binding to damaged DNA, PARP1/2 utilize NAD^+^ to add poly ADP-ribose (PAR) chains onto themselves, histones, and other protein components of the DNA repair pathway (Krishnakumar and Kraus, 2010; Messner et al., 2010; Ray Chaudhuri and Nussenzweig, 2017). These PAR chains contribute to the decompaction of chromatin and recruitment of downstream factors to coordinate the DNA damage response (DDR) (Poirier et al., 1982; Strickfaden et al., 2016). PARP1 catalyzes 85-95% of total cellular PARylation observed in response to DNA breaks (Ame et al., 1999). PARP2 which is partially redundant with PARP1 in DDR, was identified because of residual PAR activity in PARP1^-/-^cells (Ame et al., 1999; Johansson, 1999; Menissier de Murcia et al., 2003; Ronson et al., 2018). PARP1 arrives first at DNA breaks, followed by PARP2, whose recruitment is in part mediated by PARP1-dependent PARylation (Chen et al., 2018; Mahadevan et al., 2019b; Mortusewicz et al., 2007). In addition to its role in DNA repair, highly abundant PARP1 also regulates chromatin architecture and transcription (Clark et al., 2012; Kim et al., 2004; Muthurajan et al., 2014). UnPARylated PARP1 binds chromatin with high affinity and compacts it into higher order structures to block transcription *in vitro* (Kim et al., 2004; Muthurajan et al., 2014; Sukhanova et al., 2016). In addition, genomic studies have identified PARP1 at promoters of actively transcribed genes (Krishnakumar et al., 2008; Nalabothula et al., 2015).

The enzymatic activity of PARP1/2 is inhibited by a class of pharmacological agents known as PARP inhibitors (PARPi), which are NAD^+^ analogs that bind the catalytic site of PARPs to block cellular PARylation, which causes accumulation of SSBs (Rose et al., 2020; Thorsell et al., 2017). When left unrepaired in cycling cells, these SSBs are converted to DSBs that undergo homologous recombination repair (HRR) in normal cells by pathways that include the BRCA1/2 proteins. Therefore, upon treatment with PARPi, tumor cells lacking functional HRR mechanisms (i.e. BRCA^-/-^) face replication stress and cell death owing to widespread genomic instability (Farmer et al., 2005; Lord and Ashworth, 2012). This mechanism of synthetic lethality between PARPi and HRR proteins led to the recognition of PARP1/2 as key targets for cancer drugs (Bryant et al., 2005). Four PARPi (talazoparib, olaparib, niraparib and rucaparib) are now approved for treatment of breast, ovarian, and prostate cancers with HRR deficiencies (Yi et al., 2019). These and other PARPi are also being investigated for their use in combination with radiotherapy, platinum salts and other cytotoxic chemotherapeutic agents like temozolomide (Drean et al., 2016).

The physical stalling of PARP1/2 at sites of DNA damage (“PARP trapping”) has been implicated in mediating the cytotoxicity of PARPi when used in combination with alkylating agents (Blessing et al., 2020; Hopkins et al., 2015; Michelena et al., 2018; Murai et al., 2012). While it is commonly reported that various PARPi differ only marginally with respect to catalytic inhibition (IC_50_ in single digit nanomolar range), their cytotoxic potentials are vastly different. Talazoparib, the most potent PARP trapper, has the highest cytotoxic potential (Hopkins et al., 2019; Hopkins et al., 2015; Murai et al., 2014; Thorsell et al., 2017). Importantly, PARP trapping requires PARP1, but not PARP2, since only PARP1^-/-^cells have reduced sensitivity towards certain clinical PARPi stemming from loss of PARP trapping (Murai et al., 2012; Ronson et al., 2018; Shao et al., 2020).

There has been a lot of interest in elucidating the molecular mechanism of PARP trapping. *In vitro* binding assays show that PARPi do not drastically perturb the rate of release of PARP1 from DNA (Hopkins et al., 2015; Rudolph et al., 2018; Rudolph et al., 2020). While PARP1 stabilization at DNA breaks may be regulated via diverse allosteric interactions between PARPi, PARP1 and DNA substrates, these do not correlate with trapping efficiency or *in vivo* efficacy (Hopkins et al., 2015; Zandarashvili et al., 2020). The notion that PARPi physically entrap PARP1 at DNA lesions in cells has been challenged by a recent finding that PARP1 undergoes rapid turnover at DNA lesions and that PARPi do not undermine this process (Shao et al., 2020). Additionally, we have shown that talazoparib is actually much more potent than other PARPi (Rudolph et al., 2021b), suggesting that trapping potency may correlate with inhibition of activity (Rudolph et al., 2022).

A better understanding of the kinetic behavior of endogenous PARP1/2 in live cells will provide insight into the role of PARP trapping in governing the efficacy of PARPi, particularly in light of the many other roles ascribed to PARP1/2. Although the chromatin-bound and dynamic states of PARP1 have been studied using fluorescence recovery after photobleaching (FRAP), fluorescence correlation spectroscopy (FCS) and fluorescence loss in photobleaching (FLIP) in live cells transfected with fluorescently tagged expression constructs (Haince et al., 2008; Kozlowski, 2014), these bulk approaches obscure multi-state dynamic behavior. Furthermore, overexpression can potentially change the kinetic behavior of proteins (Hansen et al., 2017; Schmidt et al., 2016). Here, we set out to understand how endogenous PARP1/2 navigate the undamaged nuclear environment, move to and at DNA lesions, and then stall in the presence of PARPi. Towards this goal, we used CRISPR/Cas9-mediated genome editing to fluorescently tag endogenous PARP1/2 molecules for direct visualization using bulk and single molecule live cell microscopy. We find that PARP1/2 exist in three distinct dynamic states (fast diffusing, slow diffusing and chromatin bound) in both undamaged and damaged cells. We further categorized the chromatin bound population into transiently and stably bound PARP molecules. Upon induction of laser-induced DNA damage, only the transiently bound PARP1/2 molecules underwent stabilization whereas most molecules continued to diffuse freely. Treatment with an efficient PARP trapper, talazoparib, increased the number and retention time of stably bound PARP1 molecules at DNA lesions, but this effect did not extend to olaparib, a weaker PARP trapper. As such, our results provide key insights for the development of next-generation PARPi.

## Results

### Live-cell single molecule microscopy reveals fraction of stably bound PARP1 and PARP2 in undamaged cells

To study the dynamics of endogenous PARP1 and PARP2 molecules in live human osteosarcoma U2OS cells, we used CRISPR/Cas9 mediated genome editing to introduce sequences encoding 3X-Flag-HaloTag into the N-termini of all alleles of endogenous *parp1* or *parp2* genes. This allowed the expression of N-terminal 3X-Flag-HaloTag containing fusion proteins (Figure S1A). Accurate genome targeting was confirmed by PCR using primers flanking the two homology arms and by Sanger sequencing (Figures S1A and S1B). Robust expression of Flag-Halo-PARP1 and Flag-Halo-PARP2 was demonstrated by immunoblots (Figure S1C) and these tagged proteins were covalently modified and fluorescently labeled upon incubation with the cell permeable HaloTag ligand Janelia Fluor 646 (JF646) (Figures 1A i and S1D) (Grimm et al., 2015; Los et al., 2008).

**Figure 1.**
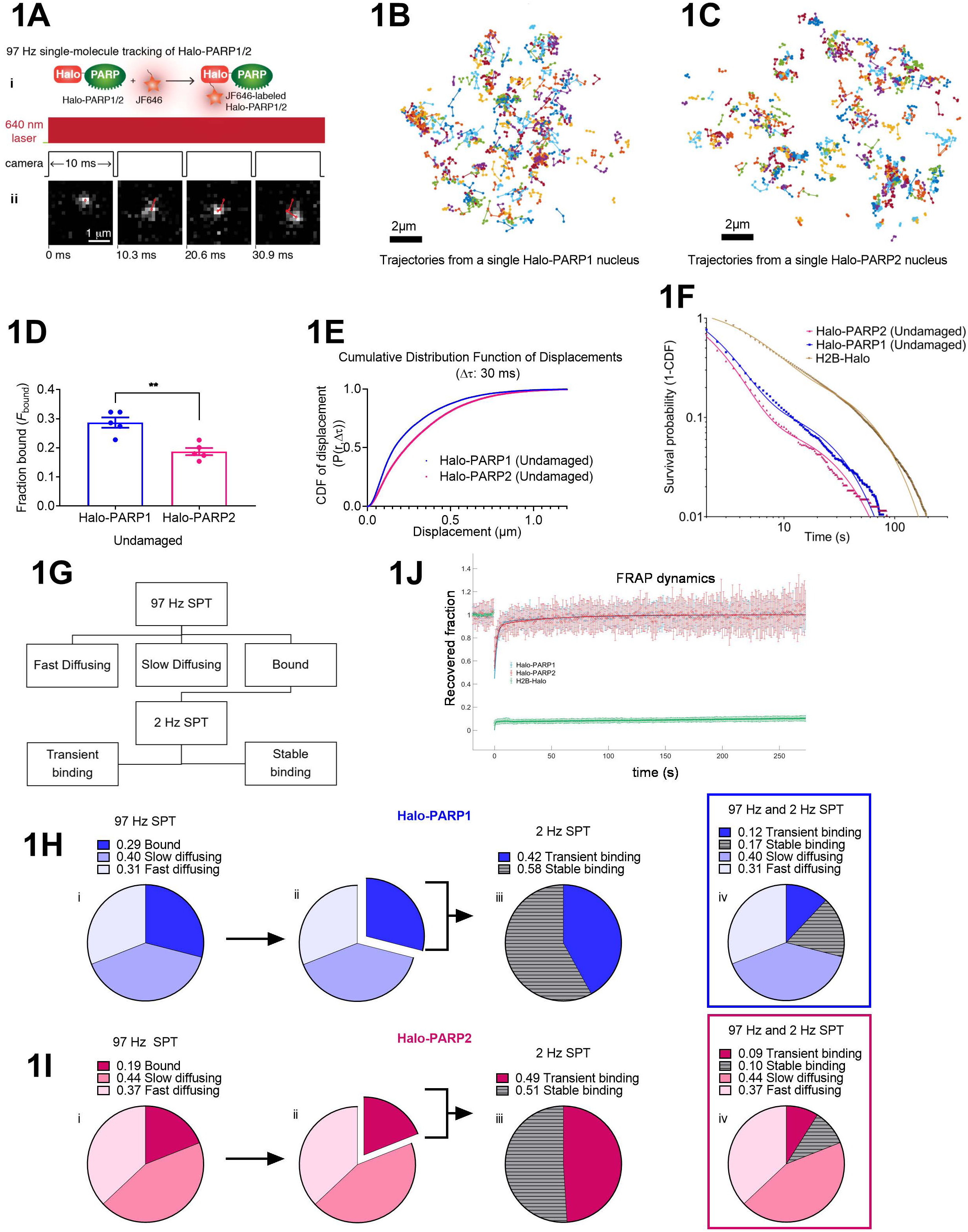
Live-cell single molecule microscopy reveals fraction of stably bound PARP1 and PARP2 in undamaged cells. **A**. i) Schematic describing the covalent binding of JF646 dye to the HaloTag. ii. Sample cropped frames from a representative 97 Hz SPT movie depicting the trajectory of a single PARP1 molecule. The 640 nm excitation laser was used continuously for imaging while the camera exposure time was 10.3 ms. **B. C**. Single particle trajectories (length of >2) over 30 s for Halo-PARP1 (in B) or Halo-PARP2 (in C) in a single representative nucleus. **D**. Fraction bound (*F*_bound_) of Halo-PARP1 and Halo-PARP2 in undamaged cells inferred from Spot-On’s three-state model fitting to 97 Hz SPT data. Bar graphs show the mean *F*_bound_ ± SEM obtained from ≥ 42,000 trajectories (>3 detections) from ≥ 52 cells from ≥ 5 independent replicates (represented by dots), each of which were fitted separately. Statistical difference between the two groups was determined using unpaired t-test. **E**. Cumulative distribution function (CDF) of displacements for Halo-PARP1 and Halo-PARP2 (representative Δ*τ* = 30 ms) in undamaged cells. Individual curves depict data merged from ≥ 42,000 trajectories (>3 detections) from ≥ 52 cells from ≥ 5 independent replicates. **F**. A log-log plot showing the uncorrected survival probability (1-CDF) of individual Halo-PARP1 and Halo-PARP2 molecules and their respective two-phase exponential model fits (solid curves) to 2 Hz SPT data in undamaged cells. Each curve represents data merged from ≥ 870 trajectories from ≥ 13 cells from ≥ 3 independent replicates. Data acquired for H2B-Halo (11,737 trajectories from ≥ 40 cells from 10 independent replicates) was used for photobleaching correction and thereby deriving values for *τ*_transient_ and *τ*_stable_ (See Table S3). **G**. Scheme showing 97 Hz and 2 Hz SPT workflow. Three-state model fits to 97 Hz SPT data using Spot-On was used to derive fractions and diffusion coefficients of fast diffusing (*F*_fast_, *D*_fast_), slow diffusing (*F*_slow_, *D*_slow_) and bound PARP (*F*_bound_, *D*_bound_) molecules. Further, 2 Hz SPT data was fit using a two-phase exponential model to derive fractions and duration of transient (Fraction transient, *τ*_transient_) and stable (Fraction stable, *τ*_stable_) PARP binding events. **H. I**. Pie chart illustrations summarizing the derivation of overall fractions of Halo-PARP1 (in H) and Halo-PARP2 (in I) engaging in transient and stable binding, slow diffusion, and fast diffusion from 97 Hz and 2 Hz SPT experiments. The bound, slow, and fast diffusing fractions (in i) were determined using Spot-On’s three-state model fitting to 97 Hz SPT data. The bound fraction (in ii) in Halo-PARP1 and Halo-PARP2 cells was analyzed by 2 Hz SPT and fit to a two-phase exponential model. Data acquired for H2B-Halo was used for photobleaching correction, and a correction factor (see Materials and Methods) was applied to obtain the true fraction of transiently and stably binding Halo-PARP molecules (in iii). These data were compiled together to obtain the overall fractions of endogenous Halo-PARP1 and Halo-PARP2 molecules (in iv). **J**. Normalized and photobleaching corrected recovery curves from FRAP experiments performed on Halo-PARP1 (blue circles) and Halo-PARP2 (red circles). H2B-Halo (green circles) was used for photobleaching correction. A two-phase exponential model (solid line) was fit to the FRAP data. Error bars represent standard deviation (SD) from 11-18 cells from ≥ 3 independent replicates.

We first validated our genome edited cell lines by performing bulk live-cell laser microirradiation (Aleksandrov et al., 2018; Mortusewicz et al., 2007) on Halo-PARP1 and Halo-PARP2 expressing U2OS cells and analyzed our data using the method of Quantitation of Fluorescence Accumulation after DNA damage (Q-FADD) (Bowerman et al., 2021; Mahadevan et al., 2019a; Mahadevan et al., 2019b). To visualize the fluorescently tagged proteins, we used a high nanomolar concentration of the HaloTag ligand, JF646. We found that endogenous Halo-PARP1 accumulates significantly faster than endogenous Halo-PARP2 at laser-induced DNA lesions, as measured by a higher effective diffusion coefficient (*D*_eff_) (Figure S1E, Table S1). This result is consistent with our previously published work using cells overexpressing GFP-PARP1 and GFP-PARP2 (Mahadevan et al., 2019b), demonstrating similar recruitment kinetics of Halo and GFP-tagged proteins. A significantly larger fraction of Halo-PARP2 was mobile (*F*_m_), compared to Halo-PARP1. This difference in *F*_m_ was revealed upon probing endogenous PARP1 and PARP2, but not with overexpressed proteins, underscoring the importance of examining the dynamics of endogenous proteins (Mahadevan et al., 2019b) (Figure S1F, Table S1).

We used our genome-edited Halo-PARP1/2 cell lines to monitor the intranuclear dynamics of individual endogenous PARP1 and PARP2 molecules in the undamaged condition. We labeled genome edited cells with a low nanomolar concentration of JF646 and visualized individual molecules of Halo-PARP1/2, as described for other Halo-tagged proteins (Grimm et al., 2015; Hansen et al., 2017; Jha and Hansen, 2022; Schmidt et al., 2016; Youmans et al., 2018). Using single particle tracking at a high frame rate (97 Hz SPT) with highly inclined and laminated optical sheet (HILO) illumination (Tokunaga et al., 2008), we could track individual PARP1/2 molecules inside the nucleus over time (Figures 1Aii, 1B and 1C). While some particles were relatively immobile and therefore presumably chromatin-bound, others displayed rapid diffusion.

To understand what fraction of PARP1/2 were bound to chromatin vs. freely diffusing, we plotted the displacement distributions of Halo-PARP1/2 and analyzed the data with ‘Spot-On’ (Hansen et al., 2018). Since a two-state model comprising bound and free fractions resulted in poor fits, especially at longer time delays Δ*τ* (Figures S1G and S1H), we instead used a three-state kinetic model consisting of bound, slow-diffusing and fast-diffusing fractions, which resulted in better fits. This suggests that in undamaged cells, endogenous PARP1/2 molecules exist in at least three distinct states: a chromatin bound state, a slow-diffusing state, and a fast-diffusing state. The latter is presumably responsible for scanning the genome for potential DNA insults. While the diffusion coefficients (for both the fast and slow diffusing fractions) are very similar between PARP1 and PARP2, a significantly higher fraction of PARP1 (0.29) than of PARP2 (0.19) exists in the chromatin bound state (*F*_bound_) (Figures 1D, 1E and Table S2). The remainder of the PARP population is distributed between the slow-diffusing fraction (*F*_slow_, PARP1 = 0.4; PARP2 = 0.44) or the fast-diffusing fraction (*F*_fast_, PARP1 = 0.31, PARP2 = 0.37) (Table S2).

Rapid photobleaching limits our ability to study stable chromatin binding events that occur at time scales longer than the 30 s length of our 97 Hz movies. We therefore adopted a different imaging scheme (2 Hz SPT) to specifically study the chromatin bound fractions of Halo-PARP1/2 using lower laser power and frame rate (2 Hz) and a longer exposure time of 500 ms over 5 min (Chen et al., 2014; Hansen et al., 2017; Huseyin and Klose, 2021). In this imaging mode, rapidly diffusing particles are blurred whereas bound molecules can be observed distinctly (Watanabe and Mitchison, 2002), thereby allowing the tracking of both stable and transient PARP binding events, as demonstrated for other nuclear proteins (Chen et al., 2014; Hansen et al., 2017; Huseyin and Klose, 2021). We studied the dissociation of Halo-PARP1/2 molecules from chromatin using 2 Hz SPT by plotting their survival probabilities. A survival curve of H2B-Halo, a histone protein known to stably associate with chromatin for multiple hours, was used as a control for photobleaching (Hansen et al., 2017; Huseyin and Klose, 2021). We fitted survival curves with a two-phase exponential decay model that accounts for binding events of PARP molecules that are either transiently or stably bound (Figures 1F and S1I). We found that a sub-fraction of 0.58 (out of 0.29 bound) PARP1 and 0.51 (out of 0.19 bound) PARP2 molecules participate in stable binding events while the remainder engages in transient binding events (Table S3).

Next, we integrated our observations from 97 Hz and 2 Hz SPT and calculated the fraction of PARP1/2 molecules that diffuse freely (either slowly or rapidly) or engage in transient and stable binding events. The resulting pie charts show that of the bound molecules, similar fractions of PARP1 and PARP2 participate in transient and stable binding events (Figures 1G-I and Table S3). Our analysis further revealed that the time constants associated with these binding events (τ_transient_ and τ_stable_) are similar for PARP1 and PARP2 in undamaged cells (Figure 1F, S1I and Table S3).

We next validated the results from 2 Hz SPT experiments using FRAP as an orthogonal approach. Upon fitting the FRAP curves of Halo-PARP1/2 with reaction-dominant model with two states (Hansen et al., 2017; Sprague et al., 2004), we found that the FRAP recovery times (PARP1 *τ*_b_ : 72.3 s; PARP2 *τ*_b_ : 58 s) are in reasonable agreement with the binding times for stable interactions (τ_stable_, PARP1: 47.6 s and PARP2: 54.7 s), as inferred from 2 Hz SPT (Figure 1J, Tables S3 and S4). Because a significantly larger fraction of PARP1 is bound to chromatin than PARP2 (from 97 Hz SPT, Table S2), it follows that the transiently and stably binding fractions of PARP1 are also larger than those of PARP2 (from 2 Hz SPT, Figures 1H and 1I).

### The majority of PARP1 and PARP2 molecules diffuse freely at laser-induced DNA lesions

To investigate how laser-induced DNA damage affects the dynamics of PARP1/2 in the nucleus, we integrated the approach of laser microirradiation with both 97 Hz and 2 Hz SPT. We first tracked PARP1/2 molecules immediately after laser induced DNA damage (405 nm) using 97 Hz SPT in a rectangular region of interest (ROI, damage region, blue) and at similar sized control regions above (red) and below the ROI (green) (Figure 2A). Upon analyzing the PARP1 data with the three-state model of Spot-On, we found that there were no significant changes in the fraction and diffusion coefficients of bound, slow, and fast diffusing PARP1 molecules at the site of laser damage, compared to unaffected areas of the nucleus (Figure 2B and Table S5). This is consistent with previous findings (Figure 1H, Tables S2 and S5) (Shao et al., 2020). In contrast, the bound fraction of PARP2 significantly increased (*F*_bound_ : from 0.2 to 0.32) at the damage region compared to other regions in the nucleus (Figure 2C, Table S5). Notably, the majority of PARP1 (0.64) and PARP2 (0.68) molecules are still not stably bound but exist in slow and fast diffusing states at the damage region (Table S5, *F*_slow_+*F*_fast_), suggesting that though DNA damage leads to an enrichment, there is still rapid exchange between the chromatin bound and freely diffusing proteins.

**Figure 2.**
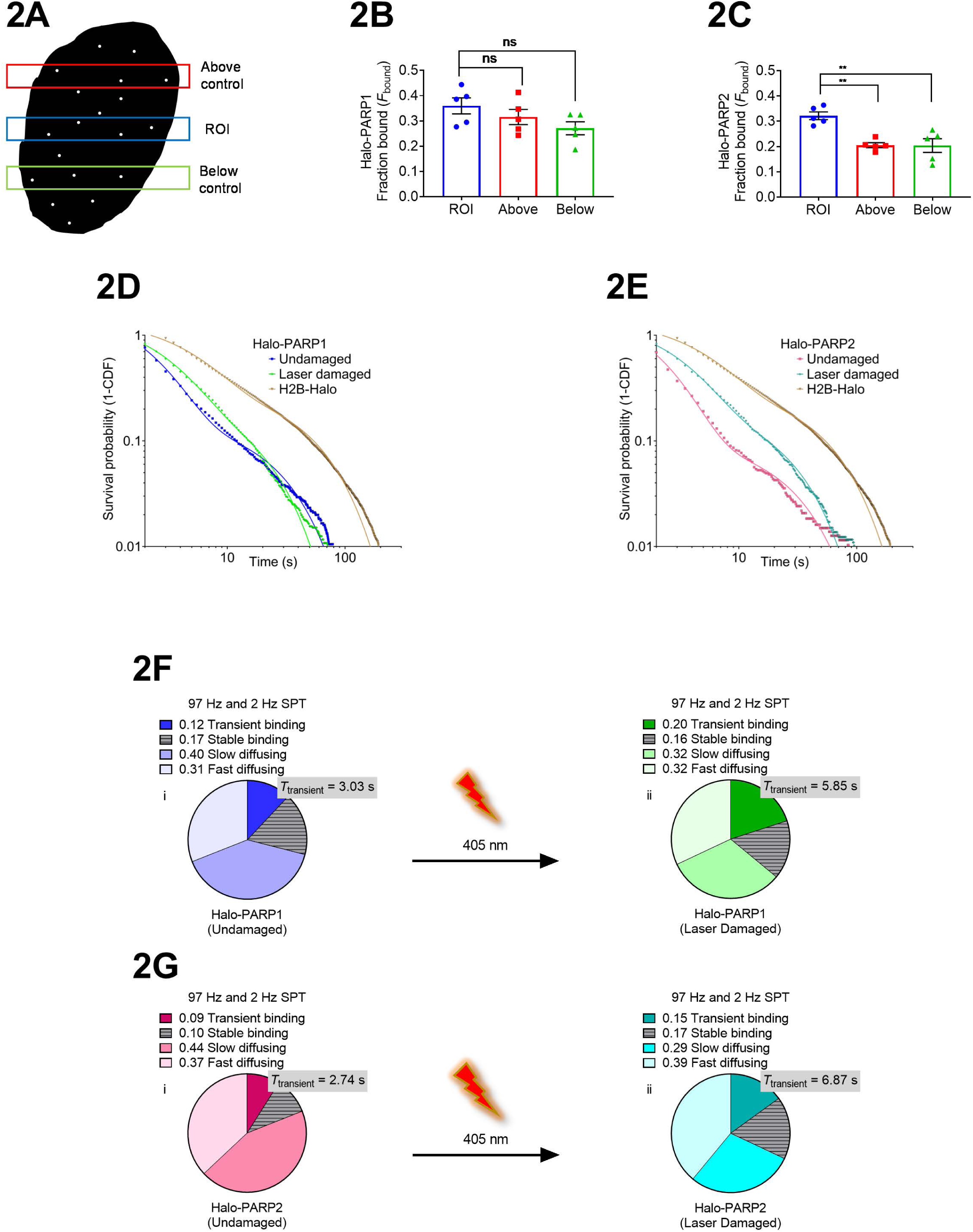
The majority of PARP1 and PARP2 molecules diffuse freely at laser-induced DNA lesions. **A**. Cartoon depicting single molecules within a nucleus and the predetermined region of interest ROI (blue), subjected to laser microirradiation, and similar sized controls above (red) and below the ROI (green). **B. C**. Fraction bound (*F*_bound_) of Halo-PARP1 (in B) and Halo-PARP2 (in C) in the ROI, above and below control regions in laser damaged cells. *F*_bound_ was inferred from Spot-On’s three-state model fitting to 97 Hz SPT data. Bar graphs show the mean *F*_bound_ ± SEM from ≥ 64,000 trajectories (>3 detections) from ≥ 64 cells from 5 independent replicates (represented by points, squares or triangles), each of which were fitted separately. Statistical differences between groups were determined using ordinary one-way ANOVA and Bonferroni’s multiple comparison tests. **D. E**. Log-log plots showing the uncorrected survival probability (1-CDF) of individual Halo-PARP1 (in D) and Halo-PARP2 (in E) molecules and their respective two-phase exponential model fits (solid curves) to 2 Hz SPT data in undamaged and laser damaged cells. Each curve represents data merged from ≥ 870 trajectories from 13-30 cells from ≥ 3 independent replicates. Data acquired for H2B-Halo (11,737 trajectories from ≥ 40 cells from 10 independent replicates) was used for photobleaching correction and thereby deriving values for *τ*_transient_ and *τ*_stable_ (See Table S3). **F. G**. Pie chart illustrations summarizing the τ_transient_ and overall fractions of Halo-PARP1 (in F) and Halo-PARP2 (in G) in undamaged (in i) and laser damaged cells (in ii). Each pie chart represents data compiled from 97 Hz and 2 Hz SPT experiments. Figures 1H iv and 1I iv were reused in 2F and 2G respectively for reference.

To better understand the properties of the fraction of PARP1 and PARP2 that are bound at regions subjected to DNA damage, we performed 2 Hz SPT and analyzed trajectories of PARP1/2 molecules immediately after the laser pulse. We found that PARP1 molecules classified in the transiently bound category were stabilized at DNA lesions compared to PARP1 molecules in undamaged cells (τ_transient_, PARP1: 3 s to 5.9 s), but still displaced ∼4 fold faster than the molecules in the stably bound category (Figures 2D, S2A and Table S6). Moreover, we saw an increase in transiently bound PARP1 molecules at the damage site (Figures 2D, S2A and Table S6). For PARP2, a similar retardation of transiently bound PARP2 molecules was observed at the damage site (τ_transient_, 2.7 s to 6.9 s), but the fraction of molecules falling into this category did not increase (Figures 2E, S2B and Table S6). Combining results from 97 Hz and 2 Hz SPT allowed us to determine how the distribution of PARP1 and PARP2 changes at DNA lesions (Figures 2F and 2G). Together, these results detail the dynamics of PARP1 and PARP2 at laser induced DNA lesions and suggest that while a majority of PARP1 and PARP2 molecules freely diffuse even in areas of intense DNA damage, the small fraction engaged in transient interactions is stabilized in areas of DNA damage, even in the absence of PARPi.

### An efficient PARP trapping agent, talazoparib, increases the retention time of only a small fraction of stably bound PARP1 molecules at damage sites

To understand how PARPi affect PARP1 exchange at laser-induced damage sites, we first performed laser microirradiation and Q-FADD analysis for bulk PARP1 accumulation in genome edited Halo-PARP1 U2OS cells treated with talazoparib, a clinical PARPi known to be the most efficient PARP trapping agent (Hopkins et al., 2019; Hopkins et al., 2015; Michelena et al., 2018; Murai et al., 2014). We observed a concentration dependent decrease in *D*_eff_ of PARP1, but not in *F*_m_, suggesting that endogenous PARP1 (when viewed as an ensemble) accumulates slower and is stalled at broken DNA ends upon treatment with talazoparib (Table S7). We followed the dynamics of bulk PARP1 release from damage foci in the presence of talazoparib. Upon fitting the portion of the curve corresponding to the decay of PARP1 with a single exponential model, we quantitated the retention time (*τ*_r_) of PARP1 at the localized damage region (Figure 3A). Talazoparib resulted in a significant concentration dependent increase in the retention time of endogenous PARP1 at chromatin regions with an abundance of DNA damage (Figure 3B, Table S7), consistent with recent findings in transfected cell lines (Blessing et al., 2020; Hendriks et al., 2021; Shao et al., 2020; Zandarashvili et al., 2020).

**Figure 3.**
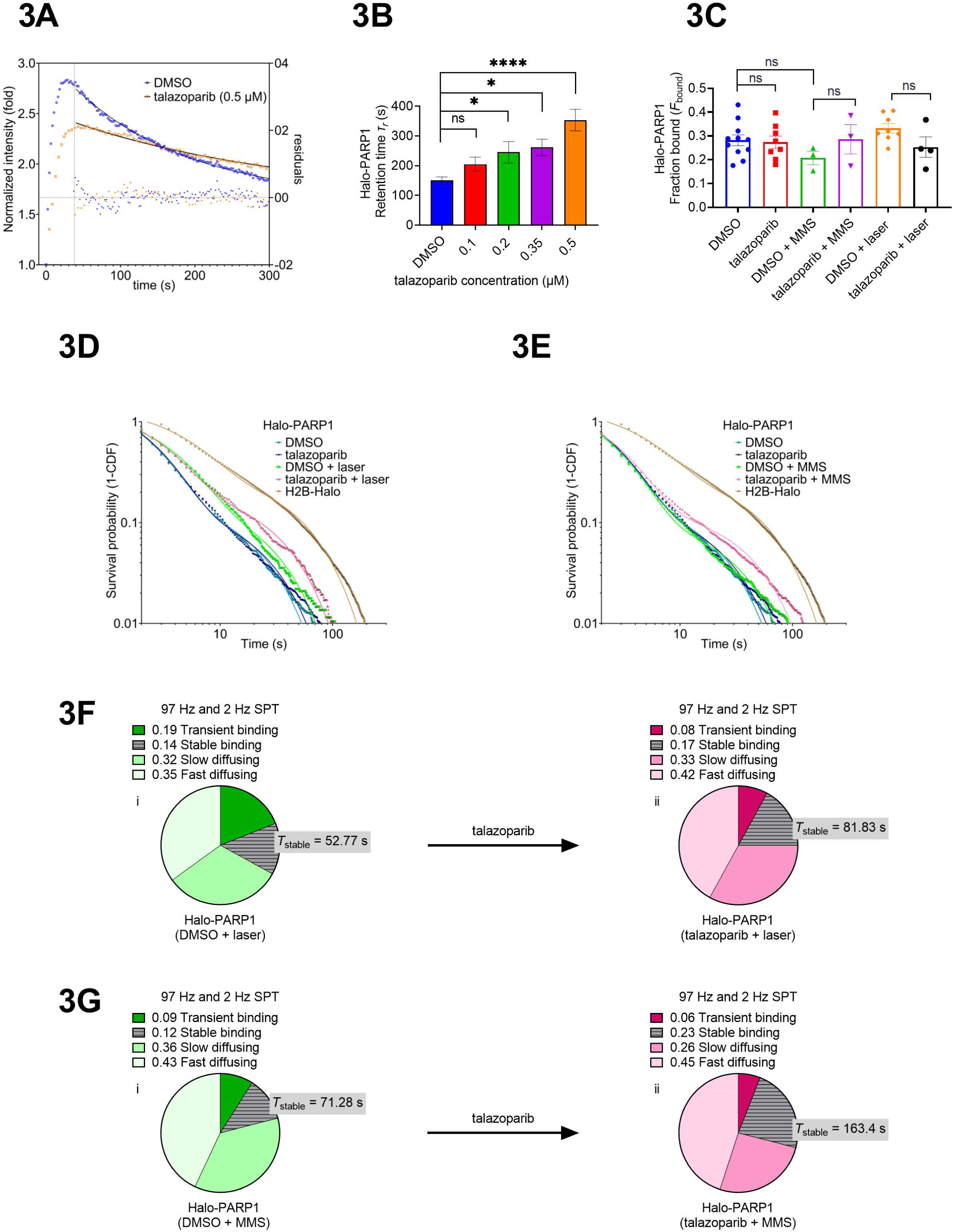
An efficient PARP trapping agent, talazoparib, increases the retention time of only a small fraction of stably binding PARP1 molecules at damage sites. **A**. Plot showing the bulk accumulation of Halo-PARP1 to and release from laser induced DNA lesions in a single representative cell treated with 0.5 μM talazoparib (orange squares) or DMSO (blue cirles). A single exponential model was fit to the portion of the kinetic curve corresponding to the release of Halo-PARP1 from DNA lesions (red curve), starting from intensity at maximum amplitude (dashed blue line). Fit residuals are shown as orange or blue dots. **B**. Halo-PARP1 retention time (*τ*_r_) derived from single exponential model fits to the portion of the kinetic curve corresponding to the release of Halo-PARP1 from DNA lesions. A plot of points representing the mean ± SEM of PARP1 retention time (*τ*_r_) from ≥ 20 cells from 2-4 independent replicate experiments/condition for increasing talazoparib concentrations (0.1 μM, 0.2 μM, 0.35 μM and 0.5 μM). Model fitting was performed individually for each cell. Statistical differences between groups were evaluated using ordinary one-way ANOVA and Bonferroni’s multiple comparisons test to compare each group with DMSO control (τ_r_ = 150.38 ± 12.17, n=28, > 3 independent replicates) **C**. Fraction bound (*F*_bound_) of Halo-PARP1 inferred from Spot-On’s three-state model fitting to 97 Hz SPT data in DMSO or talazoparib treated cells in the presence or absence of MMS or laser-induced DNA breaks. Bar graphs show the mean *F*_bound_ ± SEM from ≥ 30,000 trajectories (> 3 detections) from ≥ 45 cells from ≥ 3 independent replicates, each of which were fitted separately. Statistical differences between groups were determined using ordinary one-way ANOVA and Bonferroni’s multiple comparison tests. **D. E**. Log-log plots showing the uncorrected survival probability (1-CDF) of individual Halo-PARP1 molecules and their respective two-phase exponential model fits to 2 Hz SPT data in DMSO or talazoparib treated cells in the presence or absence of laser damage (in D) or MMS damage (in E). Each curve represents data merged from ≥ 880 trajectories from ≥ 13 cells from ≥ 3 independent replicates. Data acquired for H2B-Halo (11,737 trajectories from ≥ 40 cells from 10 independent replicates) was used for photobleaching correction and thereby deriving values for *τ*_transient_ and *τ*_stable_ (See Table S12). **F. G**. Pie chart illustrations summarizing the *τ*_stable_ and overall fractions of Halo-PARP1 in DMSO (in i) or talazoparib (in ii) treated cells in the presence of laser damage (in F) or MMS damage (in G). Each pie chart represents data compiled from 97 Hz and 2 Hz SPT experiments.

To further investigate the molecular dynamics of PARP1 stalling at DNA lesions, we utilized laser microirradiation in conjunction with 97 Hz SPT in talazoparib treated cells. With this approach, we found that talazoparib neither increased the bound fraction (*F*_bound_) nor significantly decreased the diffusion coefficients (*D*_fast_ or *D*_slow_) of PARP1 at radiation-induced DNA lesions (Figure 3C, Table S10). Similar results were also obtained when methylmethanesulfonate (MMS), an alkylating agent, was used to induce DNA damage (Figures 3C, Table S10). Together, these data imply that a majority of endogenous PARP1 molecules that were diffusing either slowly or rapidly are still doing so, and that fraction of bound PARP1 molecules didn’t change even in the presence of DNA damage and efficient PARP trapping agents.

We next performed 2 Hz SPT on Halo-PARP1 cells to characterize the bound fraction of endogenous PARP1 at DNA breaks in PARPi treated cells. We found that talazoparib increased both the fraction (0.43 to 0.70) and the duration (*τ*_stable_, 52.8 s to 81.8 s) of PARP1 molecules engaging in stable binding events and concurrently decreased the fraction (0.57 to 0.30) of transiently bound PARP1 molecules at laser induced DNA breaks (Figure 3D and S3A, Table S12). Use of MMS for DNA damage induction resulted in similar changes in the stably (0.55 to 0.79) and transiently (0.45 to 0.21) bound fractions with an even larger increase in *τ*_stable_ (71.3 s to 163.4 s) (Figures 3E, S3B and Table S12). In sum, these data suggest that talazoparib in the presence of DNA damage increases the fraction and retention time of PARP1 molecules involved in stable interactions at DNA lesions by trapping some of the transient binding PARP1 that had increased due to damage alone (Figure 2F and 2G). These results explain the slower release of PARP1 from sites of damage we observed in bulk in presence of talazoparib (Figure 3A and 3B, Table S7).

We then integrated our results from 2 Hz SPT and 97 Hz SPT experiments and deduced the overall fractions of PARP molecules participating in transient or stable binding events at laser or MMS-induced damage sites in PARPi treated cells (Figures 3F, 3G and Tables S14). These results suggest that efficient PARP trappers such as talazoparib “trap” only a small fraction of PARP1 molecules, converting some of the transient binders into stable binders at DNA lesions.

### Weaker PARP trapping agents olaparib and veliparib exert distinct effects on the retention time of stably binding PARP1 molecules

We next characterized PARP1 exchange at DNA lesions in the presence of two other well-known PARPi: olaparib (a moderate but weaker PARP1 trapping agent than talazoparib) and veliparib (a poor PARP1 trapping agent), as classified in previous studies (Hopkins et al., 2015; Murai et al., 2014). We first determined the bulk accumulation properties (*D*_eff_ and *F*_m_) and bulk retention time *τ*_r_, for Halo-PARP1 following laser induced DNA lesions upon treatment with olaparib and veliparib. At higher concentrations of olaparib, we observed a significant increase in *τ*_r_, but no concentration dependent changes were seen for values of *D*_eff_ and *F*_m_ (Figure 4A, Table S8). In contrast, treatment with increasing concentrations of veliparib did not result in significant changes in *τ*_r_, *D*_eff_ or *F*_m_ (Figure 4B, Table S9). These results suggest that both olaparib and veliparib do not impair bulk PARP1 accumulation, and that at higher concentrations, olaparib, but not veliparib, induces PARP1 stalling at DNA lesions (Figures 4A and 4B, Tables S8 and S9).

**Figure 4.**
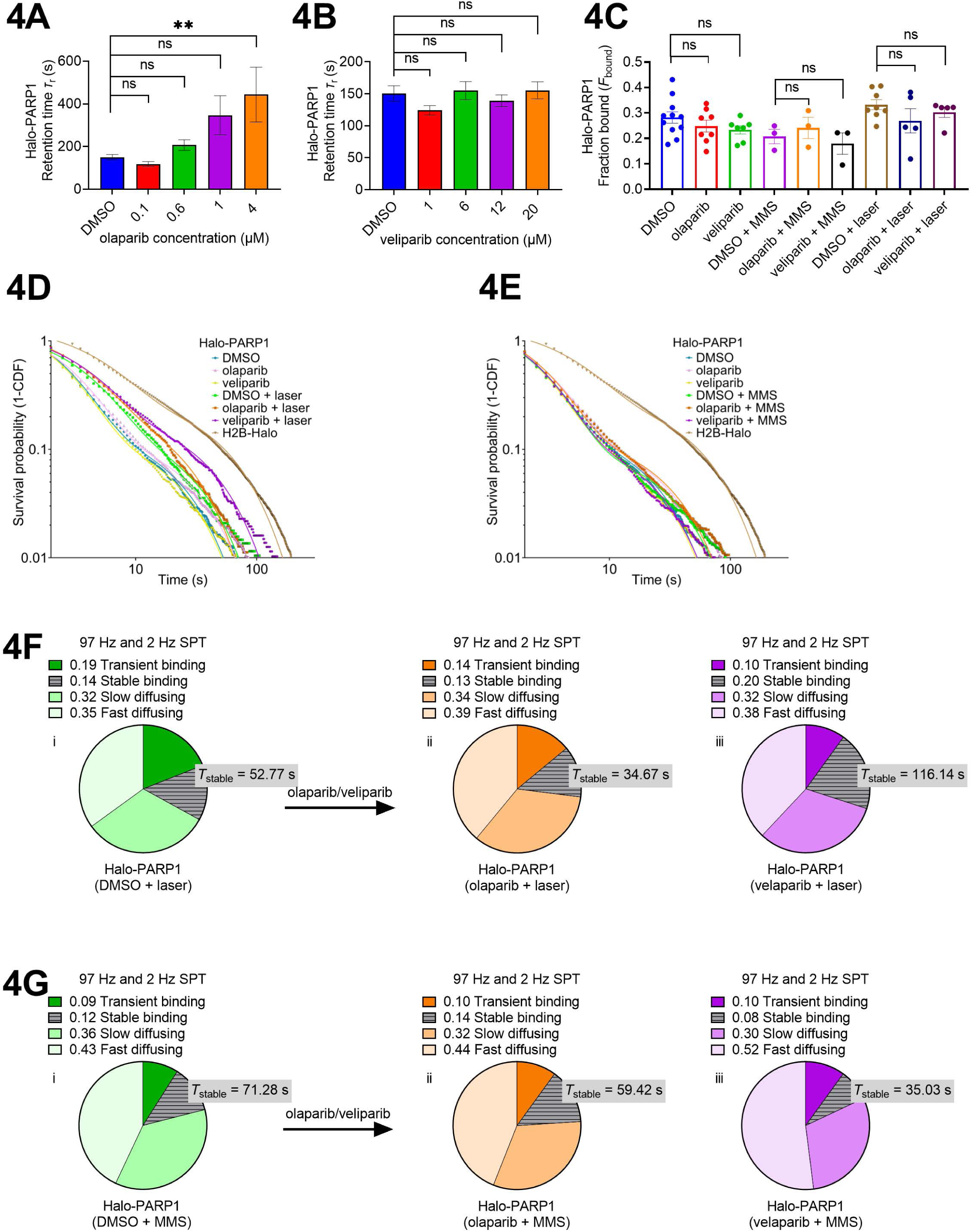
Weaker PARP trapping agents olaparib and veliparib exert distinct effects on the retention time of stably binding PARP1 molecules. **A. B**. Halo-PARP1 retention time (*τ*_r_) derived from single exponential model fits to the portion of the kinetic curve corresponding to the release of Halo-PARP1 from DNA lesions. A plot of points representing the mean ± SEM of PARP1 retention time (*τ*_r_) from ≥ 14 cells from ≥ 2 independent replicate experiments/condition for increasing olaparib concentrations (0.1 μM, 0.6 μM, 1 μM and 4 μM) (in A) and veliparib concentration (1 μM, 4 μM, 6 μM, 12 μM and 20 μM) (in B). Model fitting was performed individually for each cell. Statistical differences between groups were evaluated using ordinary one-way ANOVA and Bonferroni’s multiple comparisons test to compare each group with DMSO control (*τ*_r_ = 150.38 ± 12.17, n=28, > 3 independent replicates). **C**. Fraction bound (*F*_bound_) of Halo-PARP1 inferred from Spot-On’s three-state model fitting to 97 Hz SPT data in DMSO, olaparib and veliparib treated cells in the presence or absence of MMS or laser-induced DNA breaks. Bar graphs show the mean *F*_bound_ ± SEM from ≥ 26000 trajectories (>3 detections) from ≥ 45 cells from ≥ 3 independent replicates, each of which were fitted separately. Statistical differences between groups were determined using ordinary one-way ANOVA and Bonferroni’s multiple comparison tests. **D. E**. Log-log plots showing the uncorrected survival probability (1-CDF) of individual Halo-PARP1 molecules and their respective two-phase exponential model fits to 2 Hz SPT data in DMSO, olaparib or veliparib treated cells in the presence or absence of laser damage (in D) or MMS damage (in E). Each curve represents data merged from ≥ 874 trajectories from ≥ 12 cells from ≥ 3 independent replicates. Data acquired for H2B-Halo (11,737 trajectories from ≥ 40 cells from 10 independent replicates) was used for photobleaching correction and thereby deriving values for *τ*_transient_ and *τ*_stable_ (See Table S12). **F. G**. Pie chart illustrations summarizing the *τ*_stable_ and overall fractions of Halo-PARP1 in DMSO (in i), olaparib (in ii) or veliparib (in iii) treated cells in the presence of laser damage (in F) or MMS damage (in G). Each pie chart represents data compiled from 97 Hz and 2 Hz SPT experiments.

Upon investigating the effect of olaparib and veliparib on single molecule PARP1 dynamics at laser or MMS induced DNA breaks using 97 Hz SPT, we found that, like talazoparib, the weaker PARP trappers olaparib and veliparib did not change *F*_bound_, *D*_slow_ or *D*_fast_ of PARP1 molecules (Figure 4C, Table S10). These data suggest that neither of these drugs affects the properties of the majority of endogenous PARP1 molecules at DNA lesions. Furthermore, analysis of the bound fraction using 2 Hz SPT suggests that olaparib neither affects the binding time nor the fraction of transiently or stably interacting PARP1 molecules at laser-or MMS-induced DNA lesions (Figures 4D-E, S4A-B, Table S12). Surprisingly, at laser-induced DNA lesions, we found that veliparib had similar effects as talazoparib in increasing long-lived binding molecules (0.43 to 0.66) at the expense of more short-lived (0.57 to 0.34) PARP1 binding events. Additionally, as for treatment with talazoparib, laser damage in the presence of veliparib increased *τ*_stable_ (52.8 s to 116.1 s) of PARP1 molecules, an effect that was not observed at MMS induced DNA lesions (Figures 4D-E, S4A-B, Table S12). We then determined the overall fractions of transiently and stably binding PARP1 molecules by merging our results from 97 Hz and 2 Hz SPT for olaparib and veliparib (Figures 4F and 4G, Table S14). Collectively, these results reveal an unexpected property of veliparib in that it prolongs the dwell time of stably bound PARP1 molecules immediately after laser damage, but not after base damage induced by the alkylating agent, MMS. This is in contrast with olaparib, which does not have a significant effect on any of the properties of PARP1 in response to either type of damage.

### Trapping of stably bound PARP2 molecules is mediated by talazoparib even in the absence of DNA damage

We next applied these same methods to delineate the molecular basis of PARP2 trapping at sites of DNA lesions in PARPi treated cells. We first performed 97 Hz SPT in the presence of talazoparib, olaparib or veliparib to study PARP2 dynamics at DNA lesions induced by laser irradiation or MMS. As seen for PARP1, none of these inhibitors affected parameters of PARP2 including *F*_bound_, *D*_slow_ and *D*_fast_ at DNA lesions (Figure 5A, Table S11). Using 2 Hz SPT to study the bound fraction of PARP2 revealed that talazoparib, but not olaparib or veliparib, increased both the fraction and the duration of PARP2 molecules partaking in stable chromatin interactions both in undamaged and MMS treated cells but not in laser damaged cells (Figures 5B-C, S5A-B, Table S13). Upon integrating our results from 97 Hz and 2 Hz SPT, we determined the overall fractions of transient and stably bound PARP2 molecules in undamaged, laser damaged and MMS treated cells (Figures 5D -5F, Table S15). Together, our results suggest that entrapment of stably binding PARP2 molecules by talazoparib occurs independent of induced DNA lesions.

**Figure 5.**
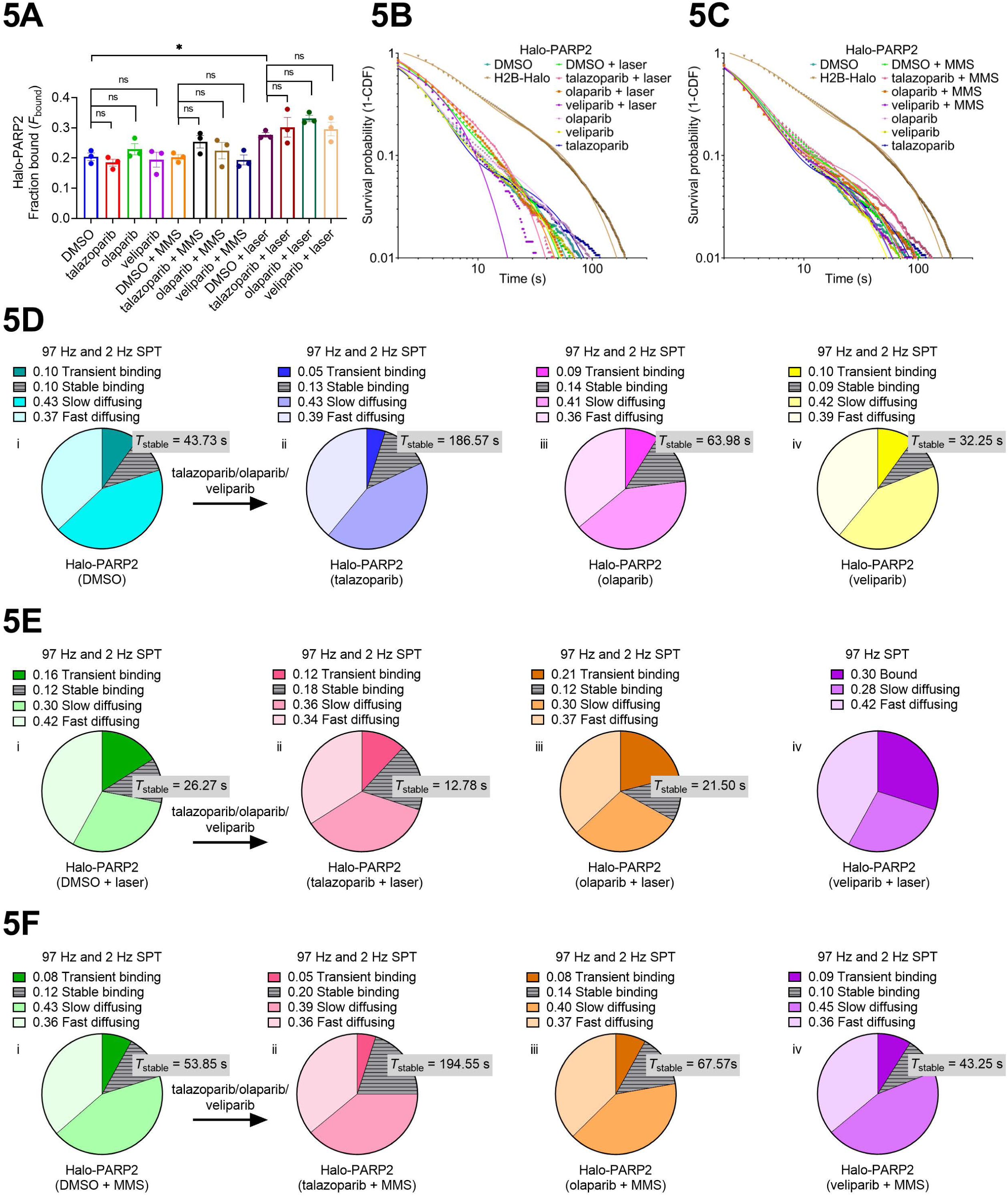
Trapping of stably bound PARP2 molecules is mediated by talazoparib even in the absence of DNA damage. **A**. Fraction bound (*F*_bound_) of Halo-PARP2 inferred from Spot-On’s three-state model fitting to 97 Hz SPT data in DMSO, olaparib and veliparib treated cells in the presence or absence of MMS or laser-induced DNA breaks. Bar graphs show the mean *F*_bound_ ± SEM from ≥ 7000 trajectories (>3 detections) from ≥ 12 cells from ≥ 3 independent replicates, each of which were fitted separately. Statistical differences between groups were determined using ordinary one-way ANOVA and Bonferroni’s multiple comparison tests. Note that we could see a significant increase in *F*_bound_ for PARP2 upon laser damage in DMSO treated cells, consistent with results shown in Figure 2C **B. C**. Log-log plots showing the uncorrected survival probability (1-CDF) of individual Halo-PARP2 molecules and their respective two-phase exponential model fits to 2 Hz SPT data in DMSO, olaparib or veliparib treated cells in the presence or absence of laser damage (in B) or MMS damage (in C). Each curve represents data merged from ≥ 625 trajectories from ≥ 12 cells from ≥ 3 independent replicates. Data acquired for H2B-Halo (11,737 trajectories from ≥ 40 cells from 10 independent replicates) was used for photobleaching correction and thereby deriving values for *τ*_transient_ and *τ*_stable_ (See Table S13). **D. E. F**. Pie chart illustrations summarizing the *τ*_stable_ and overall fractions of Halo-PARP2 in DMSO (in i), talazoparib (in ii), olaparib (in iii) or veliparib (in iv) treated cells in undamaged conditions (in D), in the presence of laser damage (in E) or MMS damage (in F). Each pie chart represents data compiled from 97 Hz and 2 Hz SPT experiments, except for veliparib in the presence of laser damage [5E (iv)], where fits to 2 Hz SPT data were unstable and inconclusive.

## Discussion

Studies spanning more than five decades have contributed to our expansive knowledge regarding the structure and biological function of PARP1 and PARP2. Here we describe for the first time, at the single-molecule level in live mammalian cells, how these abundant proteins i) navigate the native, undamaged nuclear environment, ii) recognize DNA lesions and iii) are stalled by PARPi at DNA lesions.

### Less than a third of the observed PARP1 and PARP2 population is chromatin bound in undamaged cells

PARP1 is an abundantly expressed protein with a stoichiometry of one PARP1 molecule for every ∼20 nucleosomes and is an integral component of chromatin (Kraus, 2008). Previous studies with purified protein have shown that un-PARylated PARP1 associates with intact chromatin lacking free ends (Clark et al., 2012; D’Amours et al., 1999; Muthurajan et al., 2014; Poirier et al., 1982). Genome-wide approaches have captured steady state snapshots of its genomic interactions (Krishnakumar et al., 2008; Lupey-Green et al., 2018; Nalabothula et al., 2015). *In vitro* single molecule experiments suggest that PARP1 decorates undamaged DNA and compacts it by stabilizing crossover points (Bell et al., 2021; Liu et al., 2017; Sukhanova et al., 2016). These observations collectively point to a role of PARP1 in shaping chromatin architecture. Through the direct visualization of the dynamic states of PARPs within live undamaged cells, our work revealed that less than one third of all PARP1 proteins are chromatin bound while the majority of PARP1 molecules (71%) diffuse within the nucleoplasmic space (Figure 1D, 1E and Table S2), possibly via the ‘monkey-bar’ mechanism (Rudolph et al., 2018). Even the bound fraction of PARP1 (29%) includes molecules that engage in rapid chromatin probing (transient interactions, 12% of the 29 % with *τ*_transient_ ∼3 s), and as such only a small fraction can be considered as ‘immobile’ (17% of the 29 %, *τ*_stable_ ∼48 s) (Figure 1H, Table S3).

For comparison, linker histone H1 is a major structural component of chromatin that is also abundant in cells (one H1 per nucleosome) (Bates and Thomas, 1981; Catez et al., 2006; Saha and Dalal, 2021). PARP1 and H1 reciprocally occupy gene promoters and other genomic loci (Azad et al., 2018; Krishnakumar et al., 2008). While nucleosomal core histones remain stably associated (e.g., H2B, dwell time = hours), linker histone H1 exchanges rapidly on chromatin with dwell times of only ∼3 minutes, comparable to PARP1 (∼1 min) (Table S3) (Flanagan and Brown, 2016; Lever et al., 2000; Misteli et al., 2000). PARP2, which has very different DNA-binding domains and much lower abundance than PARP1, has similar temporal characteristics as PARP1 (Figure 1I, Table S2), suggesting that its long-lived interactions may also contribute to chromatin architecture.

### Transient chromatin interactions of PARP1 and PARP2 are stabilized at laser-induced DNA lesions

Bulk laser microirradiation is one of the most widely used methods for generating localized DNA breaks to study the biological response to DNA damage in live cells (Mahadevan et al., 2019a; Zentout et al., 2021). We and others have obtained valuable insights into the recruitment of PARP1/2, demonstrating that endogenous and overexpressed PARP1 accumulates significantly faster than PARP2 at laser induced DNA lesions (Figure S1E, Table S1) (Caron et al., 2019; Haince et al., 2008; Mahadevan et al., 2019b; Mortusewicz et al., 2007). More recent work using bulk microirradiation combined with FRAP suggests that PARP1 undergoes rapid turnover at sites of DNA lesions (Shao et al., 2020). Here we have implemented a workflow that allows coupling of laser microirradiation with single particle tracking (SPT) that provide both spatial and temporal resolution for a detailed investigation of the dynamics and binding events of PARP1/2 at localized DNA lesions (Figure 2). We find that PARP1 molecules rapidly exchange at sites of DNA damage, without changes in its bound fraction (*F*_bound_) or diffusion coefficients (*D*_fast_, *D*_slow_) (Figure 2B and Table S5). Induction of DNA damage results in an increase in the local concentration of PARP1/2 (bulk accumulation of PARPs) but does not affect the behavior of the majority of proteins in damaged regions. Laser irradiation damage does increase the dwell time of the fraction of PARP1/2 molecules (for up to 7 s) partaking in transient but does not affect stable chromatin binding (Figures 2F and 2G, Table S6). Transient and stable chromatin binding modes may correspond to the different conformations of PARP1/2 on intact vs. damaged DNA. Previous studies suggest that the Zn and BRCT domains of PARP1 are involved in binding intact DNA (Rudolph et al., 2021a) whereas the Zn and WGR domains engage at broken DNA ends to activate PARP1 (Langelier et al., 2012). The transient association at DNA damage sites is sufficient for PARP1/2 to undergo self- and transPARylation (*k*_cat_ ∼ 5 – 10 s^-1^) (Langelier et al., 2008) and thus set in motion downstream repair mechanisms before they are released from DNA breaks.

### Certain PARP inhibitors trap PARP1 by extending its stable interactions with chromatin

Despite the success of PARPi in the clinic, which is attributed at least in part to ‘PARP trapping’, the molecular mechanism of PARP trapping is poorly understood. We elucidate here for the first time the changes in the dynamic properties of single particles of PARP1/2 at sites of DNA damage in the presence of PARPi. For both radiation (laser)-induced damage and chemical (MMS) damage, 97 Hz SPT revealed that the exchange of the majority of PARP1 molecules is unaffected by the presence of PARPi, irrespective of whether they are classified as efficient or inefficient PARP trapping agents (Figures 3C, 4C, Table S10). Most likely, the number of PARP1 molecules greatly exceeds the number of damage sites in our studies, and this could explain how the majority of PARP1 molecules are unaffected by DNA damage in the presence of an excess of PARPi. Our 2 Hz SPT data suggest that the effect of PARP trapping is subtle and nuanced for the different PARPi (Figures 3B and 4A, Tables S7-S9). We observe the stabilization of only the small fraction of PARP1 that participates in stable chromatin interactions. For example, for the strong PARP-trapper talazoparib, both the dwell time and the fraction (12 – 14%) of stable binding PARP1 increase upon damage induced by laser microirradiation or MMS (Figure 3F and 3G). Unexpectedly, the poor trapper veliparib shows a similar effect as talazoparib, but only after microirradiation, and not with MMS (Figure 4F, 4G). It was also unexpected that the medium trapper olaparib does not cause significant changes in fraction or dwell times for any of the populations of PARP1, bound or unbound (Figure 4F, 4G).

The subtle changes in the amount of PARP1 trapped at DNA lesions upon treatment with PARPi are in contrast to the significant fraction of apparent PARP1 trapping that is observed in bulk after cell lysis (Blessing et al., 2020; Demin et al., 2021; Hopkins et al., 2019; Hopkins et al., 2015; Michelena et al., 2018; Murai et al., 2012; Murai et al., 2014; Pommier et al., 2016). In these experiments, the overall amounts of PARP1 associated with chromatin increased with inhibitor treatment upon induction of DNA damage. In a more nuanced approach, bulk FRAP experiments showed that niraparib and talazoparib do not physically stall PARP1 at DNA lesions in live cells (Shao et al., 2020), which is consistent with our observations from single molecule experiments (Figures 3C and 4C). Our results suggest that PARP1/2 retention at DNA lesions can occur even in the absence of stable binding upon PARPi treatment. We surmise that cells treated with PARPi for extended periods of time in the presence of DNA damaging agents accumulate more and more lesions, which in turn lead to increased accumulation of PARP1 in the chromatin fraction even though the actual dynamics in the intact cell are much more subtle. It is also possible that DNA damage incurred during cell lysis and sample preparation in these previous reports yield artificially higher levels of PARP-trapping.

### Talazoparib traps stably bound PARP2 molecules independent of induced DNA breaks

PARP2 is inhibited by PARPi to the same extent as PARP1 (Rudolph et al., 2021b), an expected result given the similarity in the active sites of these two proteins. Although the efficacy of PARPi in the cell is primarily attributed to inhibition of PARP1 (Murai et al., 2012; Ronson et al., 2018; Shao et al., 2020), trapping of PARP2 at DNA lesions may contribute to the mechanism of cell toxicity. PARP2 behaves very similarly to PARP1 at sites of DNA damage wherein a majority of PARP2 molecules exchange rapidly even in the presence of PARPi (Figure 5A and Table S11). Also, as for PARP1, the bound fraction of PARP2 becomes more stable in the presence of talazoparib. For damage induced by MMS there is an increase in both fraction and dwell time whereas there is a more convoluted and subtle response for damage induced by microirradiation (Figures 5E and 5F). Interestingly, this retention effect with talazoparib is also seen in the absence of DNA damage, hinting at a special role for PARP2 in maintaining DNA integrity during replication stress (Figure 5D, Table S13). These results also point to a PARP1-activity-independent mechanism of recruitment to sites of DNA damage, one that may function in parallel or in lieu of the recently described recruitment by PARP1-mediated PARylation (Chen et al., 2018). Finally, trapping of PARP2 is not detectable with olaparib or veliparib (Figures 5D-5F, Table S13).

## Conclusion

Based on clinical experience and strong sales volume, PARPi are important and effective tools for the treatment of an increasing number of cancers. However, as with other cancer treatments that have undesired side effects (LaFargue et al., 2019) and are prone to resistance (Noordermeer and van Attikum, 2019), there is much room for improvement in the development of next-generation PARPi. Our quantitative single molecule studies in live cells challenge the classical trapping mechanism that was inferred from immunoblotting of chromatin-bound PARP1 following cell lysis (Hopkins et al., 2015; Murai et al., 2012), and independently by bulk laser microirradiation [this work and (Blessing et al., 2020; Hendriks et al., 2021; Shao et al., 2020; Zandarashvili et al., 2020)].

Our observation that the trapped fraction of PARP1 is surprisingly small suggests that more efficacious PARPi would have specificity for this small population of PARP1 that has adopted the DNA-bound conformation. Given that the concentration of PARP1 in the nucleus is many orders of magnitude larger than the amount of typical DNA damage, developing inhibitors that specifically target this minor state could reduce dosing requirements and therefore off-target effects. Promising starts in this direction have been reported (Zandarashvili et al., 2020) and we look forward to further developments in this direction.

## Limitations of the study

First, since anti-PARP1/2 antibodies failed to detect Halo-tagged PARP1/2, we could not compare the expression levels to an untagged control. We therefore cannot rule out slightly altered expression levels. Second, even though we have clearly delineated the temporal and spatial dynamics of endogenous PARP1/PARP2 in our work, the exact molecular mechanism of how slowly diffusing PARP1/2 molecules become transiently bound upon laser DNA damage, and how transiently bound PARP1/2 molecules are converted to stably bound molecules at DNA lesions in the presence of PARPi is yet to be understood. Third, as is true for all live-cell single-molecule imaging studies, although we can locate individual molecules with millisecond and nanometer precision in time and space, DNA sequence information is missing, and we do not know where in the genome PARP1/2 binds more stably, and whether these more stably bound molecules are bound to DNA damage. Future experiments may be able to overcome these limitations.

## Materials and Methods

### Mammalian cell culture

Halo-PARP1, Halo-PARP2 and parent U2OS cells were grown in McCoys 5a medium (Hyclone #SH30200) supplemented with 10% FBS, 2 mM Glutamax-I, 100 U/ml penicillin and 100 μg/mL streptomycin (complete medium). H2B-Halo-SNAP U2OS cells (kind gift from the Tjian-Darzacq laboratory, UC Berkeley, CA) were grown in low glucose DMEM medium (ThermoFisher #10567014) supplemented with 10% FBS, 2 mM Glutamax-I and 100 U/ml penicillin and 100 μg/mL streptomycin (complete medium). All cell lines used in this study were maintained in a humidified incubator at 37°C and 5% CO_2_. All cell lines were mycoplasma-free as determined by routine PCR testing.

Cells were grown and imaged on tissue culture coated CELLview slides (Greiner Bio-One # 543079) for all confocal microscopy experiments. For single molecule experiments, cells were directly grown on 35 mm circular imaging dishes (Cat # 81158) or chambered glass slides (Cat # 80807) from ibidi consisting of a #1.5H glass coverslip bottom, suitable for use in TIRF and single molecule applications.

### Endogenous tagging of parp1 and parp2 genes

To study the dynamics of endogenous PARP1 and PARP2, CRISPR/Cas9 mediated homology-directed repair (HDR) was utilized to precisely introduce HaloTag at the endogenous *parp1* and *parp2* loci with a goal of generating doubly genome edited cell lines (Xi et al., 2015). Towards this, PARP1 and PARP2 sgRNAs were inserted into the px330 plasmid (Addgene # 42230) as previously described (Cong et al., 2013). pUC19 based HDR (donor) plasmids containing the left and right homology regions were constructed using PCR and NEBuilder Hifi DNA assembly (New England Biosciences, # E2621). The template plasmid used for this process, 3xFlag-HaloTag-EZH2 HDR -pDY053, was a gift from Thomas Cech (Addgene plasmid # 171108). Briefly, 10^6^ U2OS cells were transfected with 1 μg of px330 plasmid and 1 μg of the HDR donor plasmid using the Nucleofector 2b device and cell line nucleofector kit V (Lonza, VCA-1003) per manufacturer’s protocol. Two days later, transfected cells were trypsinized and expanded in complete medium containing 1 μg/ml puromycin (ThermoFisher Scientific, # A1113803). Cells were grown in puromycin containing medium for a duration of 7 days to select cells that contain genomic integration of the HDR donor plasmid. Appropriate integration of the HaloTag was verified in the selected cell population by PCR. These cells (1.5 × 10^6^) were transfected with 2 μg of plasmid encoding eGFP-Cre recombinase (Addgene # 11923), a gift from Brian Sauer (Le et al., 1999). To obtain individual clones, cells expressing eGFP were subjected to sorting into single wells of multiple 96-well cell culture plates. Upon expansion, DNA from these cells was extracted using QuickExtract DNA extraction solution (Lucigen # QE09050) and used as a template for confirmation of homologous recombination by PCR and Sanger sequencing.

### Dye labeling

For SDS-PAGE, FRAP and bulk laser microirradiation experiments, genome edited cells were labeled with the Halo-tag ligand JF646 (a kind gift from the Lavis lab, Janelia Farms, Ashburn, VA) at a high concentration of 500 nM for 30 min at 37°C. This was followed by two washes with complete medium containing phenol red and a third wash with complete medium lacking phenol red. Cells were imaged in complete medium lacking phenol red.

For single molecule experiments, cells were labeled with JF646 at a concentration of 2 nM for both Halo-PARP1 and Halo-PARP2 cells for a duration of 30s and 2 mins respectively. For H2B-Halo-SNAP U2OS cells, JF646 was used at a concentration of 10 pM. Washes to remove extra dye were carried out as explained in the above paragraph. For all laser microirradiation experiments (bulk and single molecule), cells were sensitized to DNA damage using Hoechst 33342 (Invitrogen, Carlsbad, CA) (10 μg/mL) for 10 minutes prior to the start of imaging.

### SDS PAGE and immunoblotting

Cells were lysed using RIPA buffer (150 mM NaCl, 1.0% NP-40, 0.5% sodium deoxycholate, 0.1% SDS, 50 mM Tris, pH 8.0). The resulting whole cell protein extract was used as the protein sample for SDS PAGE and immunoblotting. The protein sample was separated on 4-12% Criterion-XT Bis-Tris gels (Bio-Rad). For SDS-PAGE experiments, JF646 fluorescence on the gel was imaged using the 647 nm channel on the Typhoon 9500 imager (GE Healthcare). For immunoblotting, monoclonal anti-FLAG M2-Peroxidase (HRP) antibody (Millipore # A8592) (1:1000) was utilized, followed by incubation with Immobilon Classico HRP substrate (# WBLUC0500). The chemiluminescence signal was detected on the Azure Biosystems GelDoc.

### PARPi and MMS treatment

All PARPi used in this study (talazoparib, olaparib and veliparib) were purchased from Selleck Chemicals and dissolved in DMSO to prepare stock solutions (2 mM). MMS (99%, Sigma Aldrich) was diluted to a concentration of 0.01 % in complete medium. Cells were treated with the indicated concentrations of PARPi and/or MMS for 1 hr at 37°C before dye-labeling and subsequent imaging. The culture medium used for dye labeling, washes and subsequent incubation during imaging contained indicated concentrations of PARPi and/or MMS.

### Live cell imaging

#### Bulk laser microirradiation

Bulk laser microirradiation was carried out as previously described (Mahadevan et al., 2019b). Briefly, a rectangular region of interest within the nucleus was subjected to DNA damage using a focused 405 nm laser beam (∼1.7 mW). Accumulation of endogenous Halo-PARP1 and Halo-PARP2 was monitored using the 647 nm laser line for 5 min.

#### Fluorescence Recovery After Photobleaching (FRAP)

FRAP experiments were performed on an inverted Nikon A1R scanning confocal microscope equipped with a 100X oil immersion objective (NA = 1.49), quad emission filter, motorized stage, 647 nm laser line and Okolab stagetop incubator for maintaining environmental conditions of temperature and humidity. Image acquisition was performed at a zoom corresponding to 256 nm x 256 nm pixel size on Nikon Elements software. A circular region of interest (radius = 10 pixels), placed away from the nuclear envelope, was bleached using the 647 nm laser line (set to 100% laser power) for 1 s. Image frames (362) were acquired at ∼2 frames per second including the first 20 pre-bleach frames for estimating initial baseline fluorescence.

#### Single molecule imaging (97 Hz and 2 Hz SPT)

Single molecule imaging experiments were carried out on a fully motorized Nikon Ti2-E inverted STORM microscope equipped with a TIRF illuminator, Agilent laser lines (405 nm, 488 nm, 561 nm and 647 nm), Nikon LU-N4 laser lines (405 nm, 488 nm, 561 nm and 640 nm) for single molecule FRAP, cage incubator for controlling temperature and humidity, two iXon Ultra 897 EMCCD cameras, 100X oil immersion TIRF objective (NA = 1.49), perfect focusing system (PFS) for correcting axial drift. These components were controlled through the NIS Elements software. All the imaging on this microscope was performed under HILO conditions wherein the incident angle was adjusted to improve the signal to background ratio (Tokunaga et al., 2008). For 97 Hz SPT experiments, images were acquired at a frame rate of ∼ 97 Hz and an exposure time of 10 ms using the 647 nm laser line (≤ 25% laser power) for a total duration of 30 s and a 128 × 128-pixel region of interest was chosen. For 2 Hz SPT experiments, images were acquired at a frame rate of 2 Hz and an exposure time of 500 ms using the 647 nm laser line set to 4% laser power for a total duration of 5 min and a 128 × 128-pixel region of interest was chosen.

### Data analysis: Bulk Laser microirradiation

Analysis of bulk microirradiation data was carried out using qFADD.py. qFADD.py is a Python implementation of the Q-FADD algorithm and its preprocessing steps, that includes the improvements of correction for nuclear drift and automated grid-search for identifying the best-fit model (Bowerman et al., 2021). The source code for qFADD.py is available at https://github.com/Luger-Lab/Q-FADD. Bulk dissipation kinetics from the DNA damage region were determined from the ensemble of individual dissipation trajectories, each fit using a single-exponential model. The reported value for retention time is the average across the ensemble of all probed nuclei. Treating each trajectory as an individual datapoint of the population, rather than averaging the dissipation trajectories, allows us to account for the effects of individual nuclear shapes on the underlying kinetics (Mahadevan et al., 2019b) and to determine the error in the ensemble metric by evaluating the standard error of the mean retention time across all nuclei within an experimental condition (i.e., PARPi type and concentration).

### Data analysis: Fluorescence Recovery After Photobleaching (FRAP)

FRAP data was analyzed using a custom-written image analysis pipeline as previously described (Hansen et al., 2017). Briefly, movies were read in, the nucleus was segmented by thresholding after the application of a gaussian filter. Fluorescence intensity in the whole nucleus and in the bleach spot was then quantified over time and background corrected. We used the total nuclear intensity to normalize for photobleaching. We manually corrected for drift. After these corrections, FRAP recovery curves from individual nuclei were averaged to obtain a mean recovery curve. To extract a residence time, we fit a reaction-dominant two-state exponential model (Sprague et al., 2004) to the FRAP curve:

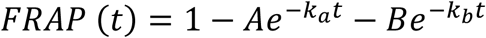

where *k*_a_ and *k*_b_ are the faster and slower off-rates respectively. Dwell times for transiently (*τ*_a_) and stably binding (*τ*_b_) molecules were calculated as follows:

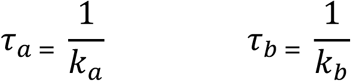

### Data analysis: Localization and tracking of SPT movies

All movies obtained from single molecule imaging were processed using a custom MATLAB implementation of the ‘multiple target tracing (MTT)’ algorithm (Hansen et al., 2017; Serge et al., 2008). This implementation is available on GitLab: https://gitlab.com/tjian-darzacq-lab/SPT_LocAndTrack

The following parameters were used to process 97 Hz SPT movies: localization error = 10^−6.25^, number of deflation loops = 0, number of gaps allowed in trajectories = 1, maximum expected diffusion coefficient = 6 μm^2^/s. The following parameters were used to process 2 Hz SPT movies: localization error = 10^−6.25^, number of deflation loops = 0, number of gaps allowed in trajectories = 2, maximum expected diffusion coefficient = 0.25 μm^2^/s. Additional parameters including distance of control ROIs (above and below) from the damage ROI = 3 μm and number of additional pixels on all sides for uniform expansion = 3 were used for processing SPT + laser microirradiation data.

### Data analysis: Analysis of trajectories from fast 97 Hz SPT movies

To analyze trajectories from the fast 97 Hz movies, we used Spot-On (Hansen et al., 2018). Spot-On performs kinetic modeling of displacements to extract the fraction and diffusion coefficient of each subpopulation. Briefly, in Spot-On we model diffusion as Brownian and model particles as existing in either a bound state (low diffusion coefficient) or one or more diffusive states. Since state transitions are neglible at the fast frame rate of 97 Hz, they are not modeled (Hansen et al., 2018). A major bias in the analysis of fast 97 Hz SPT data is defocalization. Since we are performing 2D imaging of a 3D nucleus, molecules can move out of focus axially and the rate of defocalization depends strongly on the diffusion coefficient. Spot-On corrects explicitly for this, by modeling loss due to axial diffusion over time and leverages the rate of defocalization as additional information to constrain the estimation of the diffusion coefficient.

We found that a 3-state model consisting of a bound, a slowly diffusing, and a fast-diffusing subpopulation was necessary to fit our SPT data. Thus, the distribution of displacements was fit to:

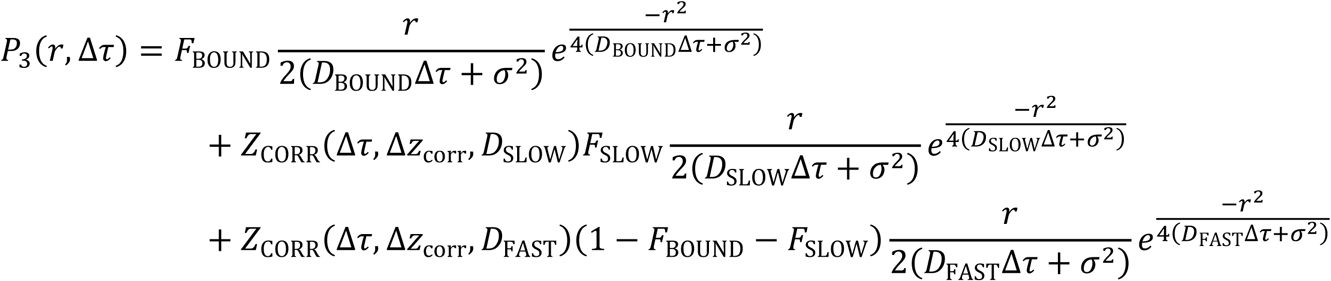

where:

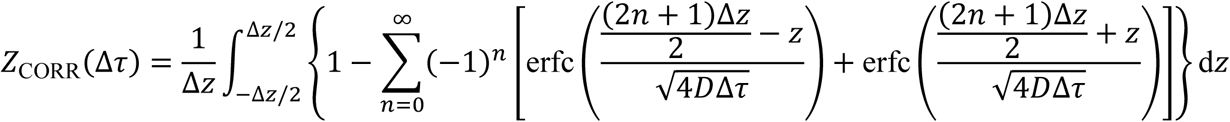

and:

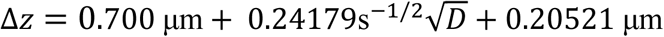

Here, *F*^BOUND^ is the fraction of molecules that are bound to chromatin, *D*^BOUND^ is the diffusion constant of chromatin bound molecules, *D*^SLOW^ is diffusion constant of the slow subpopulation of freely diffusing molecules, *D*FAST is the diffusion constant of the slow subpopulation of freely diffusing molecules, *r* is the displacement length, Δ*τ* is lag time between frames, Δ*z* is axial detection range, *σ* is localization error and *Z*CORR corrects for defocalization bias (i.e. the fact that freely diffusing molecules gradually move out-of-focus, but chromatin bound molecules do not).

Model fitting for 97 Hz SPT movies was done using Spot-On’s three state model to derive diffusion coefficient and fraction of fast diffusing, slow diffusing and bound molecules (Hansen et al., 2018). The following input parameters were used for this analysis: kinetic model = 3 state, *D*_bound_ = 0.0005 -0.08 μm^2^/s, *D*_slow_ = 0.15 -5 μm^2^/s, *D*_fast_ = 0.5 -25 μm^2^/s, *F*_bound_ and *F*_fast_ = 0 – 1, localization error = 0.048, dZ = 0.7 μm, Model fit = CDF (Cumulative distribution function) and iterations = 3. The code is freely available at https://gitlab.com/tjian-darzacq-lab/Spot-On-cli

### Data analysis: Analysis of trajectories from slow 2 Hz SPT movies

Data obtained from 2 Hz SPT experiments were used to plot merged survival curves (from multiple cells imaged over ≥ 3 independent replicates; 99% of all trajectories were used for analysis) indicating the survival probability (1-CDF) of particles at a given time (s). We fitted a two-phase exponential model (GraphPad Prism) to these survival curves to derive the fraction and dwell time (*τ*) of transient and stable binding events. The following constraints were placed on parameters: Plateau = 0, τ_transient_ and τ_stable_ > 0.

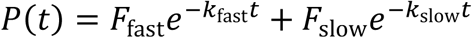

Similarly, a double-exponential was fit to the survival curve of H2B and the slow component was then used to correct for photobleaching as previously described (Hansen et al., 2017) according to:

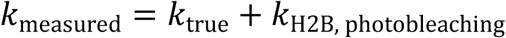

This allows us to extract the photobleaching-corrected residence time according to:

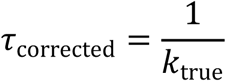

Due to the transient binding events being shorter than the stable binding events, they will inherently be overcounted. For example, suppose you have a transient residence time of 1 sec and a stable residence time of 100 sec, where the ON and OFF rates are identical (and *F*fast = *F*slow). During a 200 sec observation window, even though the same number of proteins will be stably and transiently bound, we will observe 100 transient binding events for every stable binding event. Thus, to correct for this bias, we used the following formula:

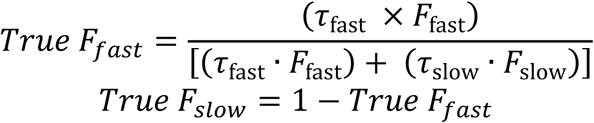

### Statistical analysis

Statistical testing for all experiments was conducted using GraphPad Prism 9. For every experiment, 2 -10 independent replications were performed. Two-tailed, unpaired Student’s t-tests were used to determine statistical significance between two groups of data. For experiments with > 2 groups of data, ordinary one-way ANOVA and Bonferroni’s multiple comparison’s tests were used to test for statistical significance. Levels of statistical significance were defined as follows: ns (not significant) p > 0.05, * p < 0.05, ** p < 0.01, *** p < 0.001, **** p < 0.0001. We have included in the figure and table legends, details of the number of cells used, the number of independent replicates, data representation in the form of merged data, mean ± SEM or mean ± SD, statistical tests used and the significance levels. If no significance levels are indicated in tables, it implies ns (p > 0.05)

For 2 Hz SPT experiments, survival curves consisting of merged data from multiple cells imaged over ≥ 3 independent replicates were plotted and compared. Owing to the sparse population of bound PARP1 and PARP2 molecules, determination of the mean ± SEM was not possible for these datasets. As a solution to this problem, we identified merged datasets with the highest number of PARP1 molecules and used their independent replicates to determine mean ± SEM. This exercise was performed for multiple merged PARP1 datasets, including undamaged, laser damaged and PARPi treated datasets. Only if the percent difference between two experimental groups was greater than the determined percent SEM for that parameter, was it considered to be a statistically significant difference (indicated by #)

## Acknowledgements

We thank Dr. Luke Lavis (Howard Hughes Medical Institute Janelia Research Campus, Ashburn, VA) for generously sharing the HaloTag JF646 dye. We also thank Uma Muthurajan (University of Colorado Boulder), Guillaume Gaullier (Uppsala University, Sweden), Daniel Youmans (Anschutz Medical Campus, CO) and Jens Schmidt (Michigan State University, MI) for helpful discussions and suggestions, Joe Dragavon (University of Colorado Boulder BioFrontiers Advanced Light Microscopy Core, RRID: SCR_018302) for help with imaging and Theresa Nahreini (University of Colorado Boulder Biochemistry Cell Culture Facility; S10ODO21601) for assistance with flow cytometry. Laser scanning confocal microscopy was performed on a Nikon A1R microscope supported by NIST-CU Cooperative Agreement award number 70NANB15H226. Single molecule microscopy was performed on a Nikon Ti-E microscope supported by the Howard Hughes Medical Institute. J.M. is a recipient of a postdoctoral fellowship award 20POST35211059 from the American Heart Association. Funding for this work was also provided by the National Cancer Institute (R01 CA218255 to K.L.) and by the Howard Hughes Medical Institute (to K.L.). A.S.H. gratefully acknowledges funding from the NIH (R00GM130896, DP2GM140938, R33CA257878, UM1HG011536, NSF (2036037), Mathers’ Foundation, and a Pew-Stewart Cancer Research Scholar grant.

---

**Table S1.** Mean effective diffusion coefficients (*D*_eff_) and Fraction mobile (*F*_m_) ± SEM for Halo-PARP1 and Halo-PARP2 inferred from Q-FADD analysis performed on individual cells subjected to bulk laser microirradiation. Statistical difference between the two groups was determined using unpaired t-tests. (Related to Figure 1)

|  | Halo-PARP1 | Halo-PARP2 |
| --- | --- | --- |
| $D_{\text{eff}}$ ($\mu\text{m}^2/\text{s}$) | $3.52 \pm 0.32$ | $0.24 \pm 0.04^{****}$ |
| $F_m$ | $0.4 \pm 0.02$ | $0.53 \pm 0.03^{****}$ |

**Table S2.** Mean diffusion coefficients and fractions ± SEM for bound, slow, and fast molecules of Halo-PARP1 and Halo-PARP2 in undamaged cells. Statistical difference between the two groups was determined using unpaired t-tests. (Related to Figure 1)

| | Diffusion coefficient ($\mu\text{m}^2/\text{s}$) | | Fraction | |
| --- | --- | --- | --- | --- |
|  | Halo-PARP1 | Halo-PARP2 | Halo-PARP1 | Halo-PARP2 |
| **Bound** | $0.01 \pm 0.002$ | $0.01 \pm 0.004$ | $0.29 \pm 0.02$ | $0.19 \pm 0.01^{**}$ |
| **Slow** | $0.58 \pm 0.06$ | $0.74 \pm 0.15$ | $0.4 \pm 0.04$ | $0.44 \pm 0.03$ |
| **Fast** | $2.88 \pm 0.24$ | $3.34 \pm 0.65$ | $0.31 \pm 0.05$ | $0.37 \pm 0.03$ |

**Table S3.** Photobleaching corrected *τ* and true fraction of transiently and stably binding Halo-PARP1 and Halo-PARP2 molecules in undamaged cells, derived from two-phase exponential model fits to 2 Hz SPT data. For details on testing for statistical significance, refer to the section of ‘Statistical analysis’ in Materials and Methods. (Related to Figure 1)

| Two-phase exponential model | Halo-PARP1 (undamaged) | Halo-PARP2 (undamaged) |
| --- | --- | --- |
| **Fraction transient** | 0.42 | 0.49 |
| **Fraction stable** | 0.58 | 0.51 |
| $\tau_{\text{transient}}$ (s) | 3.03 | 2.74 |
| $\tau_{\text{stable}}$ (s) | 47.62 | 54.67 |

**Table S4.** List of key parameters inferred from two-phase exponential fits to Halo-PARP1 and Halo-PARP2 merge FRAP data (Related to Figure S1)

| FRAP |  |  |
| --- | --- | --- |
| Two-phase exponential model<br>$1-A \cdot \exp(-k_a \cdot t) - B \cdot \exp(-k_b \cdot t)$ | | |
| Photobleaching corrected | Halo-PARP1 | Halo-PARP2 |
| **A** | 0.49 | 0.36 |
| **$\tau_A$ (s)** | 2.54 | 2.24 |
| **B** | 0.06 | 0.08 |
| **$\tau_B$ (s)** | 72.3 | 58.06 |

**Table S5.**
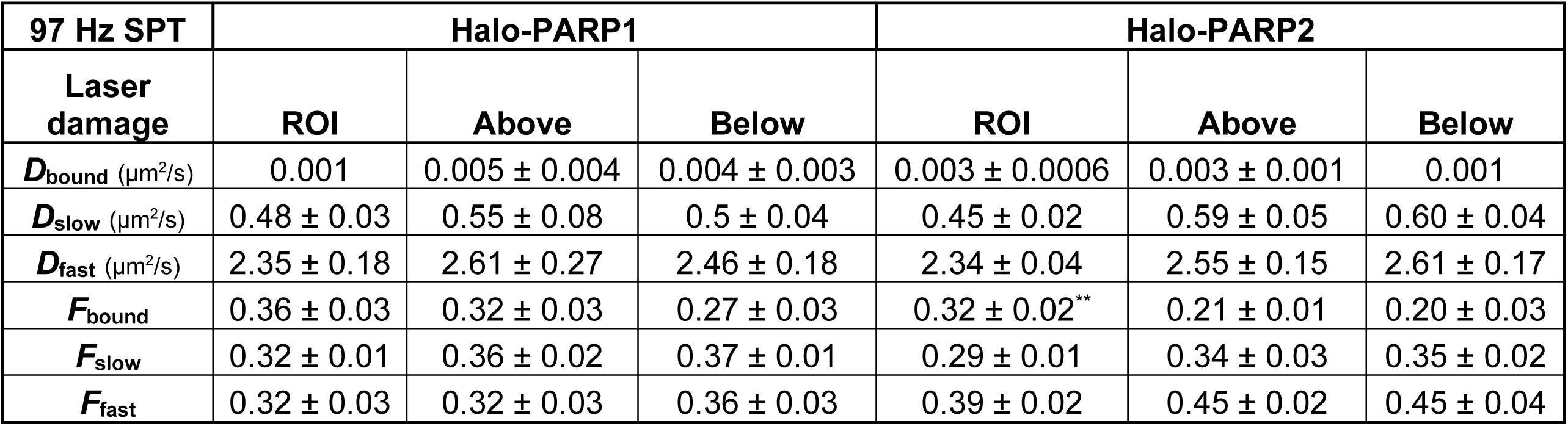
A table of key parameters (mean ± SEM) inferred from Spot-On’s three-state model fitting to 97 Hz SPT data in laser-induced DNA lesions (ROI) or control regions (Above or Below) in Halo-PARP1 and Halo-PARP2 cells. Statistical differences between groups were determined using ordinary one-way ANOVA and Bonferroni’s multiple comparison tests between ROI and above or below controls regions (Related to Figure 2)

**Table S6.** Photobleaching corrected *τ* and true fraction of transiently and stably binding Halo-PARP1 and Halo-PARP2 molecules in undamaged and laser damaged cells, derived from two-phase exponential model fits to 2 Hz SPT data. For details on testing for statistical significance, refer to the section of ‘Statistical analysis’ in Materials and Methods. (Related to Figure 2)

| Two-phase exponential model | Halo-PARP1 (undamaged) | Halo-PARP1 (laser damaged) | Halo-PARP2 (undamaged) | Halo-PARP2 (laser damaged) |
| --- | --- | --- | --- | --- |
| **Fraction transient** | 0.42 | 0.55<sup>#</sup> | 0.49 | 0.48 |
| **Fraction stable** | 0.58 | 0.45<sup>#</sup> | 0.51 | 0.52 |
| **$\tau_{\text{transient}}$ (s)** | 3.03 | 5.85<sup>#</sup> | 2.74 | 6.87<sup>#</sup> |
| **$\tau_{\text{stable}}$ (s)** | 47.62 | 23.4 | 54.67 | 42.96 |

**S7. Related to Figure 3.** Mean ± SEM of retention time *τ*_r_ derived from single exponential model fits to Halo-PARP1 release and *D*_eff_ and *F*_m_ derived from Q-FADD analysis performed on Halo-PARP1 accumulation in bulk laser microirradiation experiments upon treatment with increasing concentration of talazoparib (in S7), olaparib (in S8) and veliparib (in S9). Statistical differences between groups were evaluated using ordinary one-way ANOVA and Bonferroni’s multiple comparisons test to compare each group with DMSO control (τ_r_ = 150.38 ± 12.17, D_eff_ = 2.345 ± 0.2, F_m_ = 0.41 ± 0.02)

| Halo-PARP1 | talazoparib |  |  |
| --- | --- | --- | --- |
| Concentrations ($\mu\text{M}$) | $\tau_r$ (s) | $D_{\text{eff}}$ ($\mu\text{m}^2/\text{s}$) | $F_m$ |
| **0.1** | 205.07 ± 23.65 | 2.07 ± 0.23 | 0.33 ± 0.02 * |
| **0.2** | 245.41 ± 36.17 * | 1.86 ± 0.18 | 0.29 ± 0.02 **** |
| **0.35** | 261.60 ± 27.84 * | 1.90 ± 0.13 | 0.45 ± 0.02 |
| **0.5** | 353.75 ± 36.11 **** | 1.50 ± 0.11 | 0.43 ± 0.02 |

**S8. Related to Figure 4.**
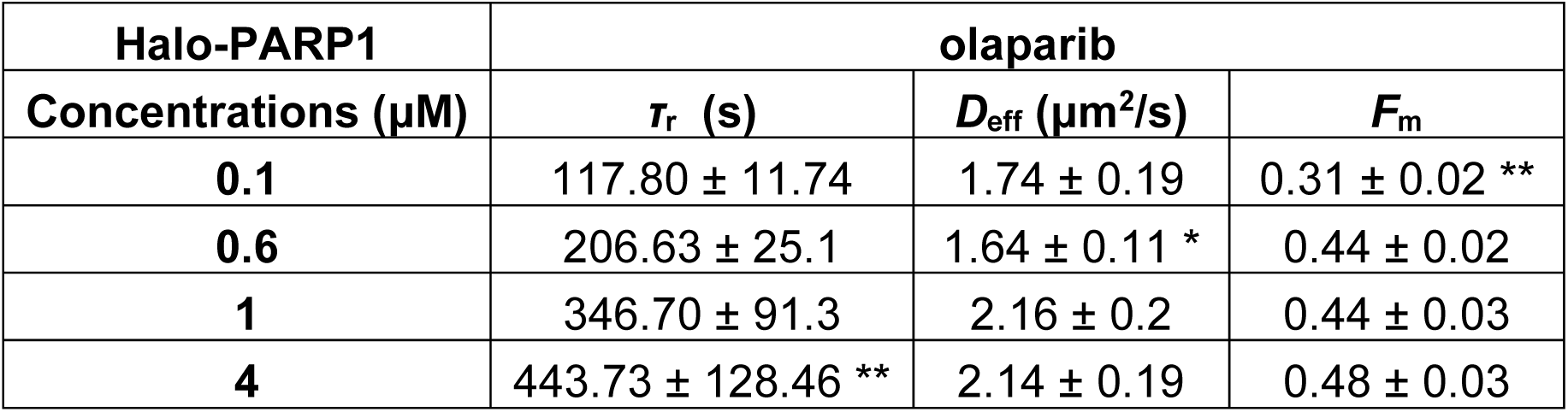
Mean ± SEM of retention time *τ*_r_ derived from single exponential model fits to Halo-PARP1 release and *D*_eff_ and *F*_m_ derived from Q-FADD analysis performed on Halo-PARP1 accumulation in bulk laser microirradiation experiments upon treatment with increasing concentration of talazoparib (in S7), olaparib (in S8) and veliparib (in S9). Statistical differences between groups were evaluated using ordinary one-way ANOVA and Bonferroni’s multiple comparisons test to compare each group with DMSO control (τ_r_ = 150.38 ± 12.17, D_eff_ = 2.345 ± 0.2, F_m_ = 0.41 ± 0.02)

| Halo-PARP1 | olaparib |  |  |
| --- | --- | --- | --- |
| Concentrations ($\mu\text{M}$) | $\tau_r$ (s) | $D_{\text{eff}}$ ($\mu\text{m}^2/\text{s}$) | $F_m$ |
| **0.1** | 117.80 ± 11.74 | 1.74 ± 0.19 | 0.31 ± 0.02 ** |
| **0.6** | 206.63 ± 25.1 | 1.64 ± 0.11 * | 0.44 ± 0.02 |
| **1** | 346.70 ± 91.3 | 2.16 ± 0.2 | 0.44 ± 0.03 |
| **4** | 443.73 ± 128.46 ** | 2.14 ± 0.19 | 0.48 ± 0.03 |

**S9. Related to Figure 4.** Mean ± SEM of retention time *τ*_r_ derived from single exponential model fits to Halo-PARP1 release and *D*_eff_ and *F*_m_ derived from Q-FADD analysis performed on Halo-PARP1 accumulation in bulk laser microirradiation experiments upon treatment with increasing concentration of talazoparib (in S7), olaparib (in S8) and veliparib (in S9). Statistical differences between groups were evaluated using ordinary one-way ANOVA and Bonferroni’s multiple comparisons test to compare each group with DMSO control (τ_r_ = 150.38 ± 12.17, D_eff_ = 2.345 ± 0.2, F_m_ = 0.41 ± 0.02)

| Halo-PARP1 | veliparib |  |  |
| --- | --- | --- | --- |
| Concentrations ($\mu\text{M}$) | $\tau_r$ (s) | $D_{\text{eff}}$ ($\mu\text{m}^2/\text{s}$) | $F_m$ |
| **1** | 124.23 ± 7.28 | 2.64 ± 0.2 | 0.43 ± 0.02 |
| **4** | 106.77 ± 7.14 | 2.45 ± 0.35 | 0.33 ± 0.02 * |
| **6** | 154.92 ± 14.15 | 2.20 ± 0.19 | 0.42 ± 0.02 |
| **12** | 139.14 ± 9.1 | 1.70 ± 0.11 | 0.40 ± 0.02 |
| **20** | 155.46 ± 13.34 | 2.30 ± 0.23 | 0.38 ± 0.02 |

**S10.** Tables listing key parameters (mean ± SEM) inferred from Spot-On’s three-state model fitting to 97 Hz SPT data in DMSO, talazoparib, olaparib and veliparib treated cells in the presence or absence of laser or MMS induced DNA lesions in Halo-PARP1 (in S10) and Halo-PARP2 (in S11) cells. Statistical differences between groups were determined using ordinary one-way ANOVA and Bonferroni’s multiple comparison tests of drug treated groups with their respective vehicle treated groups. (Related to Figures 3, 4 and 5)

**S10.**
|  | Halo-PARP1 |  |  |  |  |  |
| --- | --- | --- | --- | --- | --- | --- |
| 97 Hz SPT | $D_{\text{bound}}$ ($\mu\text{m}^2/\text{s}$) | $D_{\text{slow}}$ ($\mu\text{m}^2/\text{s}$) | $D_{\text{fast}}$ ($\mu\text{m}^2/\text{s}$) | $F_{\text{bound}}$ | $F_{\text{slow}}$ | $F_{\text{fast}}$ |
| DMSO | 0.006 ± 0.001 | 0.7 ± 0.02 | 3.23 ± 0.15 | 0.28 ± 0.02 | 0.35 ± 0.02 | 0.37 ± 0.03 |
| talazoparib | 0.007 ± 0.002 | 0.76 ± 0.06 | 3.49 ± 0.19 | 0.27 ± 0.03 | 0.35 ± 0.01 | 0.38 ± 0.03 |
| olaparib | 0.013 ± 0.005 | 0.7 ± 0.05 | 3.2 ± 0.12 | 0.25 ± 0.02 | 0.34 ± 0.02 | 0.41 ± 0.02 |
| veliparib | 0.008 ± 0.003 | 0.7 ± 0.03 | 3.12 ± 0.15 | 0.23 ± 0.02 | 0.34 ± 0.01 | 0.43 ± 0.03 |
| DMSO + laser | 0.005 ± 0.002 | 0.56 ± 0.03 | 2.82 ± 0.1 | 0.33 ± 0.02 | 0.32 ± 0.01 | 0.35 ± 0.01 |
| talazoparib + laser | 0.005 ± 0.001 | 0.64 ± 0.07 | 3.04 ± 0.23 | 0.25 ± 0.04 | 0.33 ± 0.02 | 0.42 ± 0.04 |
| olaparib + laser | 0.008 ± 0.004 | 0.69 ± 0.09 | 3.11 ± 0.19 | 0.27 ± 0.05 | 0.34 ± 0.04 | 0.39 ± 0.04 |
| veliparib + laser | 0.004 ± 0.001 | 0.64 ± 0.03 | 3.07 ± 0.07 | 0.30 ± 0.02 | 0.32 ± 0.01 | 0.38 ± 0.02 |
| DMSO + MMS | 0.008 ± 0.005 | 0.55 ± 0.09 | 2.5 ± 0.20 | 0.21 ± 0.03 | 0.36 ± 0.05 | 0.43 ± 0.08 |
| talazoparib + MMS | 0.006 ± 0.003 | 0.55 ± 0.03 | 2.70 ± 0.13 | 0.29 ± 0.06 | 0.26 ± 0.02 | 0.45 ± 0.07 |
| olaparib + MMS | 0.01 ± 0.007 | 0.54 ± 0.08 | 2.67 ± 0.2 | 0.24 ± 0.04 | 0.32 ± 0.03 | 0.44 ± 0.06 |
| veliparib + MMS | 0.009 ± 0.003 | 0.646 ± 0.05 | 2.77 ± 0.17 | 0.18 ± 0.04 | 0.30 ± 0.02 | 0.52 ± 0.06 |

**S11.** Tables listing key parameters (mean ± SEM) inferred from Spot-On’s three-state model fitting to 97 Hz SPT data in DMSO, talazoparib, olaparib and veliparib treated cells in the presence or absence of laser or MMS induced DNA lesions in Halo-PARP1 (in S10) and Halo-PARP2 (in S11) cells. Statistical differences between groups were determined using ordinary one-way ANOVA and Bonferroni’s multiple comparison tests of drug treated groups with their respective vehicle treated groups. (Related to Figures 3, 4 and 5)

**S11.**
|  | Halo-PARP2 |  |  |  |  |  |
| --- | --- | --- | --- | --- | --- | --- |
| 97 Hz SPT | $D_{\text{bound}}$ ($\mu\text{m}^2/\text{s}$) | $D_{\text{slow}}$ ($\mu\text{m}^2/\text{s}$) | $D_{\text{fast}}$ ($\mu\text{m}^2/\text{s}$) | $F_{\text{bound}}$ | $F_{\text{slow}}$ | $F_{\text{fast}}$ |
| DMSO | 0.02 ± 0.005 | 0.73 ± 0.05 | 2.97 ± 0.11 | 0.20 ± 0.01 | 0.43 ± 0.03 | 0.37 ± 0.03 |
| talazoparib | 0.02 ± 0.001 | 0.96 ± 0.03 * | 4.02 ± 0.09 * | 0.18 ± 0.01 | 0.43 ± 0.03 | 0.39 ± 0.04 |
| olaparib | 0.01 ± 0.003 | 0.84 ± 0.06 | 3.72 ± 0.31 | 0.23 ± 0.02 | 0.41 ± 0.01 | 0.36 ± 0.02 |
| veliparib | 0.02 ± 0.001 | 0.97 ± 0.04 * | 3.9 ± 0.19 * | 0.19 ± 0.02 | 0.42 ± 0.04 | 0.39 ± 0.06 |
| DMSO + laser | 0.002 ± 0.001 | 0.62 ± 0.15 | 2.73 ± 0.51 | 0.28 ± 0.01 * | 0.3 ± 0.06 | 0.42 ± 0.06 |
| talazoparib + laser | 0.009 ± 0.004 | 0.63 ± 0.06 | 3.08 ± 0.11 | 0.3 ± 0.03 | 0.36 ± 0.02 | 0.34 ± 0.05 |
| olaparib + laser | 0.004 ± 0.001 | 0.61 ± 0.04 | 3.01 ± 0.22 | 0.33 ± 0.01 | 0.30 ± 0.03 | 0.37 ± 0.03 |
| veliparib + laser | 0.007 ± 0.003 | 0.54 ± 0.11 | 2.784 ± 0.35 | 0.30 ± 0.02 | 0.28 ± 0.01 | 0.42 ± 0.02 |
| DMSO + MMS | 0.02 ± 0.001 | 0.79 ± 0.06 | 3.41 ± 0.27 | 0.20 ± 0.01 | 0.43 ± 0.02 | 0.36 ± 0.02 |
| talazoparib + MMS | 0.01 ± 0.003 | 0.91 ± 0.01 | 4.17 ± 0.38 | 0.25 ± 0.02 | 0.39 ± 0.02 | 0.36 ± 0.04 |
| olaparib + MMS | 0.02 ± 0.001 | 0.97 ± 0.05 | 4.46 ± 0.17 | 0.22 ± 0.03 | 0.40 ± 0.004 | 0.37 ± 0.03 |
| veliparib + MMS | 0.02 ± 0.002 | 0.94 ± 0.06 | 4.12 ± 0.27 | 0.19 ± 0.02 | 0.45 ± 0.002 | 0.36 ± 0.02 |

**S12.**
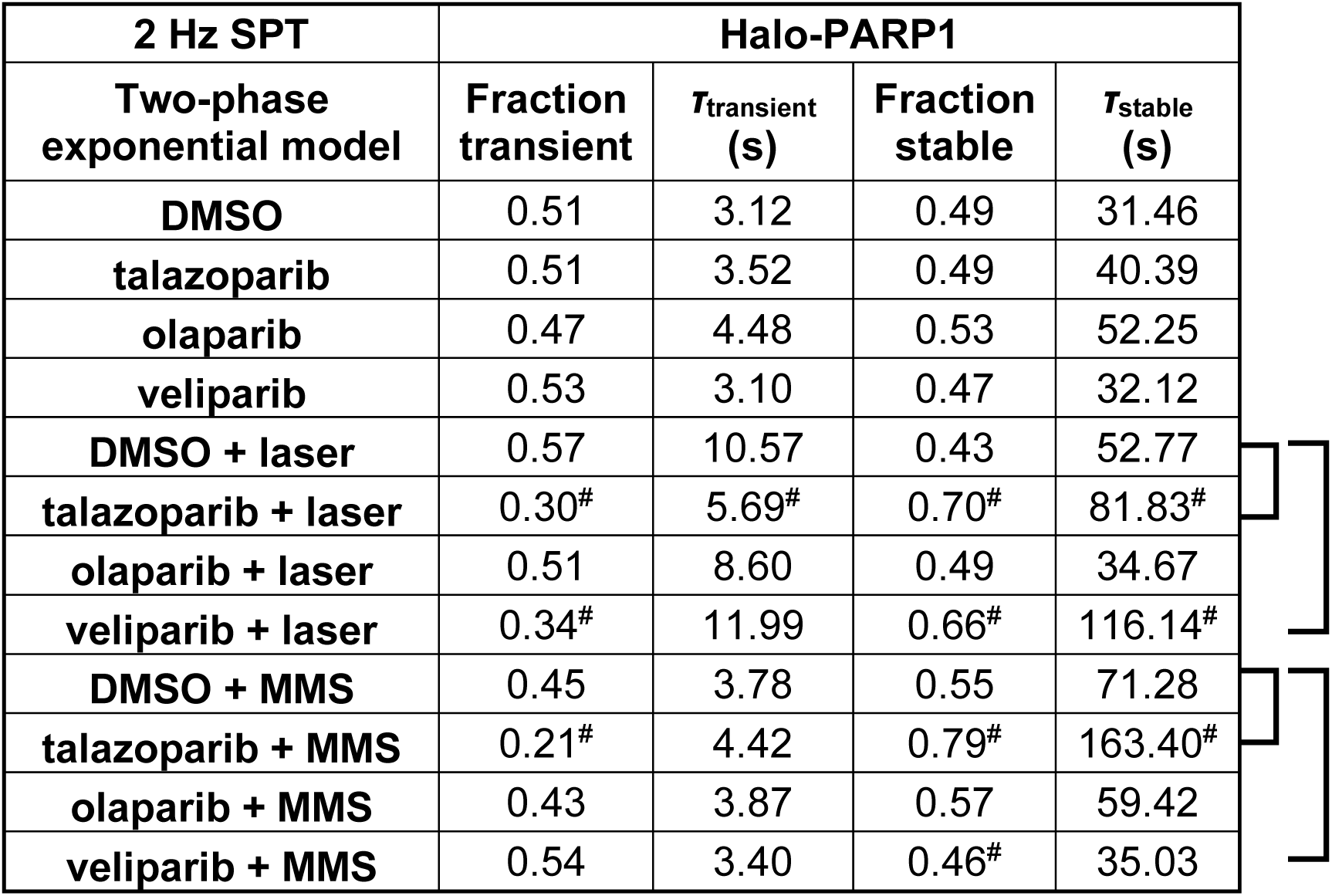
Photobleaching corrected *τ* and true fraction of transiently and stably binding Halo-PARP1 (in S12) and Halo-PARP2 (in S13) molecules, derived from two-phase exponential model fits to 2 Hz SPT data in DMSO, talazoparib, olaparib and veliparib treated cells in the presence or absence of laser or MMS induced DNA lesions. For details on testing for statistical significance, refer to the section of ‘Statistical analysis’ in Materials and Methods. (Related to Figures 3, 4 and 5)

**S12.**
| 2 Hz SPT | Halo-PARP1 |  |  |  |
| --- | --- | --- | --- | --- |
| Two-phase exponential model | Fraction transient | $\tau_{\text{transient}}$ (s) | Fraction stable | $\tau_{\text{stable}}$ (s) |
| DMSO | 0.51 | 3.12 | 0.49 | 31.46 |
| talazoparib | 0.51 | 3.52 | 0.49 | 40.39 |
| olaparib | 0.47 | 4.48 | 0.53 | 52.25 |
| veliparib | 0.53 | 3.10 | 0.47 | 32.12 |
| DMSO + laser | 0.57 | 10.57 | 0.43 | 52.77 |
| talazoparib + laser | 0.30<sup>#</sup> | 5.69<sup>#</sup> | 0.70<sup>#</sup> | 81.83<sup>#</sup> |
| olaparib + laser | 0.51 | 8.60 | 0.49 | 34.67 |
| veliparib + laser | 0.34<sup>#</sup> | 11.99 | 0.66<sup>#</sup> | 116.14<sup>#</sup> |
| DMSO + MMS | 0.45 | 3.78 | 0.55 | 71.28 |
| talazoparib + MMS | 0.21<sup>#</sup> | 4.42 | 0.79<sup>#</sup> | 163.40<sup>#</sup> |
| olaparib + MMS | 0.43 | 3.87 | 0.57 | 59.42 |
| veliparib + MMS | 0.54 | 3.40 | 0.46<sup>#</sup> | 35.03 |

**S13.** Photobleaching corrected *τ* and true fraction of transiently and stably binding Halo-PARP1 (in S12) and Halo-PARP2 (in S13) molecules, derived from two-phase exponential model fits to 2 Hz SPT data in DMSO, talazoparib, olaparib and veliparib treated cells in the presence or absence of laser or MMS induced DNA lesions. For details on testing for statistical significance, refer to the section of ‘Statistical analysis’ in Materials and Methods. (Related to Figures 3, 4 and 5)

**S13**
| 2 Hz SPT | Halo-PARP2 |  |  |  |
| --- | --- | --- | --- | --- |
| Two-phase exponential model | Fraction transient | $\tau_{\text{transient}}$ (s) | Fraction stable | $\tau_{\text{stable}}$ (s) |
| DMSO | 0.51 | 3.62 | 0.49 | 43.73 |
| talazoparib | 0.27<sup>#</sup> | 4.14 | 0.73<sup>#</sup> | 186.57<sup>#</sup> |
| olaparib | 0.41 | 3.85 | 0.59 | 63.98 |
| veliparib | 0.52 | 3.20 | 0.48 | 34.25 |
| DMSO + laser | 0.57 | 5.09 | 0.43 | 26.27 |
| talazoparib + laser | 0.41 | 6.77 | 0.59 | 12.78 |
| olaparib + laser | 0.63 | 3.33 | 0.37 | 21.50 |
| veliparib + laser | Not determined : Unstable fits |  |  |  |
| DMSO + MMS | 0.42 | 3.81 | 0.58 | 53.85 |
| talazoparib + MMS | 0.21<sup>#</sup> | 4.55 | 0.79<sup>#</sup> | 194.55<sup>#</sup> |
| olaparib + MMS | 0.36 | 2.80 | 0.64 | 67.57 |
| veliparib + MMS | 0.48 | 3.07 | 0.52 | 43.25 |

**S14.** Lists summarizing the overall true fractions of transiently binding, stably binding, slow and fast diffusing Halo-PARP1 (in S14) and Halo-PARP2 (in S15) derived from analysis of 97 Hz and 2 Hz SPT data in DMSO, talazoparib, olaparib and veliparib treated cells in the presence or absence of laser or MMS induced DNA lesions. (Related to Figures 3, 4 and 5)

**S14.**
| **Halo-PARP1** | **$F_{\text{bound}}$** |  |  |  |
| --- | --- | --- | --- | --- |
| **97 Hz and 2 Hz SPT** | **Fraction transient** | **Fraction stable** | **$F_{\text{slow}}$** | **$F_{\text{fast}}$** |
| **DMSO** | 0.14 | 0.14 | 0.35 | 0.37 |
| **talazoparib** | 0.14 | 0.13 | 0.35 | 0.38 |
| **olaparib** | 0.12 | 0.13 | 0.34 | 0.41 |
| **veliparib** | 0.12 | 0.11 | 0.34 | 0.43 |
| **DMSO + laser** | 0.19 | 0.14 | 0.32 | 0.35 |
| **talazoparib + laser** | 0.075 | 0.175 | 0.33 | 0.42 |
| **olaparib + laser** | 0.14 | 0.13 | 0.34 | 0.39 |
| **veliparib + laser** | 0.10 | 0.20 | 0.32 | 0.38 |
| **DMSO + MMS** | 0.09 | 0.12 | 0.36 | 0.43 |
| **talazoparib + MMS** | 0.06 | 0.23 | 0.26 | 0.45 |
| **olaparib + MMS** | 0.10 | 0.14 | 0.32 | 0.44 |
| **veliparib + MMS** | 0.10 | 0.08 | 0.30 | 0.52 |

**S15.**
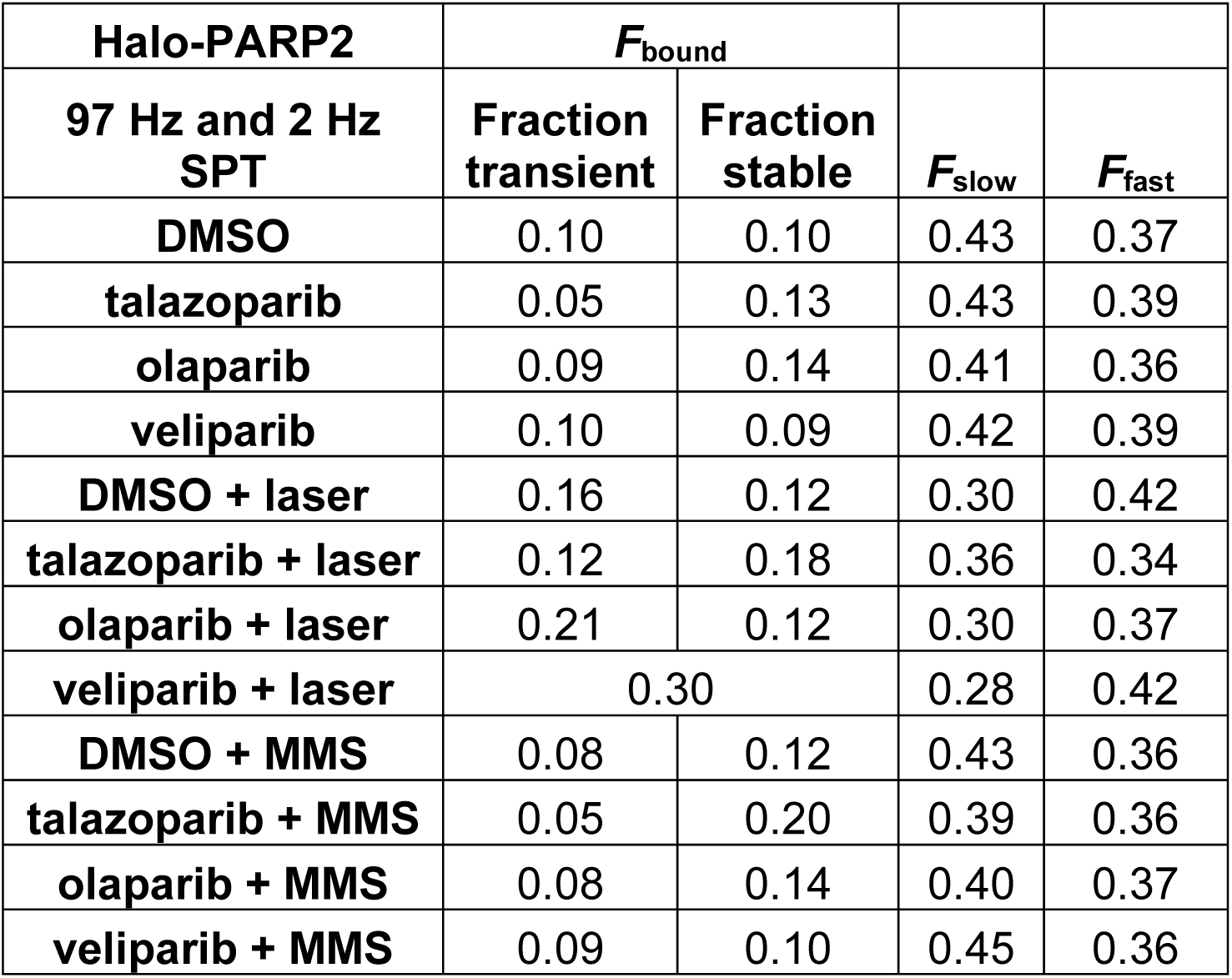
Lists summarizing the overall true fractions of transiently binding, stably binding, slow and fast diffusing Halo-PARP1 (in S14) and Halo-PARP2 (in S15) derived from analysis of 97 Hz and 2 Hz SPT data in DMSO, talazoparib, olaparib and veliparib treated cells in the presence or absence of laser or MMS induced DNA lesions. (Related to Figures 3, 4 and 5)

## References

Aleksandrov, R., Dotchev, A., Poser, I., Krastev, D., Georgiev, G., Panova, G., Babukov, Y., Danovski, G., Dyankova, T., Hubatsch, L., et al. (2018). Protein Dynamics in Complex DNA Lesions. Mol Cell 69, 1046–1061 e1045.

Ame, J.C., Rolli, V., Schreiber, V., Niedergang, C., Apiou, F., Decker, P., Muller, S., Hoger, T., Menissier-de Murcia, J., and de Murcia, G. (1999). PARP-2, A novel mammalian DNA damage-dependent poly(ADP-ribose) polymerase. J Biol Chem 274, 17860–17868.

Azad, G.K., Ito, K., Sailaja, B.S., Biran, A., Nissim-Rafinia, M., Yamada, Y., Brown, D.T., Takizawa, T., and Meshorer, E. (2018). PARP1-dependent eviction of the linker histone H1 mediates immediate early gene expression during neuronal activation. J Cell Biol 217, 473–481.

Bates, D.L., and Thomas, J.O. (1981). Histones H1 and H5: one or two molecules per nucleosome? Nucleic Acids Res 9, 5883–5894.

Bell, N.A.W., Haynes, P.J., Brunner, K., de Oliveira, T.M., Flocco, M.M., Hoogenboom, B.W., and Molloy, J.E. (2021). Single-molecule measurements reveal that PARP1 condenses DNA by loop stabilization. Sci Adv 7.

Benjamin, R.C., and Gill, D.M. (1980). Poly(ADP-ribose) synthesis in vitro programmed by damaged DNA. A comparison of DNA molecules containing different types of strand breaks. J Biol Chem 255, 10502–10508.

Blessing, C., Mandemaker, I.K., Gonzalez-Leal, C., Preisser, J., Schomburg, A., and Ladurner, A.G. (2020). The Oncogenic Helicase ALC1 Regulates PARP Inhibitor Potency by Trapping PARP2 at DNA Breaks. Mol Cell 80, 862–875 e866.

Bowerman, S., Mahadevan, J., Benson, P., Rudolph, J., and Luger, K. (2021). Automated Modeling of Protein Accumulation at DNA Damage Sites using qFADD.py. bioRxiv, 2021.2003.2015.435501.

Bryant, H.E., Schultz, N., Thomas, H.D., Parker, K.M., Flower, D., Lopez, E., Kyle, S., Meuth, M., Curtin, N.J., and Helleday, T. (2005). Specific killing of BRCA2-deficient tumours with inhibitors of poly(ADP-ribose) polymerase. Nature 434, 913–917.

Caron, M.C., Sharma, A.K., O’Sullivan, J., Myler, L.R., Ferreira, M.T., Rodrigue, A., Coulombe, Y., Ethier, C., Gagne, J.P., Langelier, M.F., et al. (2019). Poly(ADP-ribose) polymerase-1 antagonizes DNA resection at double-strand breaks. Nat Commun 10, 2954.

Catez, F., Ueda, T., and Bustin, M. (2006). Determinants of histone H1 mobility and chromatin binding in living cells. Nat Struct Mol Biol 13, 305–310.

Chen, J., Zhang, Z., Li, L., Chen, B.C., Revyakin, A., Hajj, B., Legant, W., Dahan, M., Lionnet, T., Betzig, E., et al. (2014). Single-molecule dynamics of enhanceosome assembly in embryonic stem cells. Cell 156, 1274–1285.

Chen, Q., Kassab, M.A., Dantzer, F., and Yu, X. (2018). PARP2 mediates branched poly ADP-ribosylation in response to DNA damage. Nat Commun 9, 3233.

Clark, N.J., Kramer, M., Muthurajan, U.M., and Luger, K. (2012). Alternative modes of binding of poly(ADP-ribose) polymerase 1 to free DNA and nucleosomes. J Biol Chem 287, 32430–32439.

Cong, L., Ran, F.A., Cox, D., Lin, S., Barretto, R., Habib, N., Hsu, P.D., Wu, X., Jiang, W., Marraffini, L.A., et al. (2013). Multiplex genome engineering using CRISPR/Cas systems. Science 339, 819–823.

D’Amours, D., Desnoyers, S., D’Silva, I., and Poirier, G.G. (1999). Poly(ADP-ribosyl)ation reactions in the regulation of nuclear functions. Biochem J 342 (Pt 2), 249–268.

Demin, A.A., Hirota, K., Tsuda, M., Adamowicz, M., Hailstone, R., Brazina, J., Gittens, W., Kalasova, I., Shao, Z., Zha, S., et al. (2021). XRCC1 prevents toxic PARP1 trapping during DNA base excision repair. Mol Cell 81, 3018–3030 e3015.

Drean, A., Lord, C.J., and Ashworth, A. (2016). PARP inhibitor combination therapy. Crit Rev Oncol Hematol 108, 73–85.

Farmer, H., McCabe, N., Lord, C.J., Tutt, A.N., Johnson, D.A., Richardson, T.B., Santarosa, M., Dillon, K.J., Hickson, I., Knights, C., et al. (2005). Targeting the DNA repair defect in BRCA mutant cells as a therapeutic strategy. Nature 434, 917–921.

Flanagan, T.W., and Brown, D.T. (2016). Molecular dynamics of histone H1. Biochim Biophys Acta 1859, 468–475.

Grimm, J.B., English, B.P., Chen, J., Slaughter, J.P., Zhang, Z., Revyakin, A., Patel, R., Macklin, J.J., Normanno, D., Singer, R.H., et al. (2015). A general method to improve fluorophores for live-cell and single-molecule microscopy. Nat Methods 12, 244–250, 243 p following 250.

Haince, J.F., McDonald, D., Rodrigue, A., Dery, U., Masson, J.Y., Hendzel, M.J., and Poirier, G.G. (2008). PARP1-dependent kinetics of recruitment of MRE11 and NBS1 proteins to multiple DNA damage sites. J Biol Chem 283, 1197–1208.

Hansen, A.S., Pustova, I., Cattoglio, C., Tjian, R., and Darzacq, X. (2017). CTCF and cohesin regulate chromatin loop stability with distinct dynamics. Elife 6.

Hansen, A.S., Woringer, M., Grimm, J.B., Lavis, L.D., Tjian, R., and Darzacq, X. (2018). Robust model-based analysis of single-particle tracking experiments with Spot-On. Elife 7.

Hendriks, I.A., Buch-Larsen, S.C., Prokhorova, E., Elsborg, J.D., Rebak, A., Zhu, K., Ahel, D., Lukas, C., Ahel, I., and Nielsen, M.L. (2021). The regulatory landscape of the human HPF1-and ARH3-dependent ADP-ribosylome. Nat Commun 12, 5893.

Hopkins, T.A., Ainsworth, W.B., Ellis, P.A., Donawho, C.K., DiGiammarino, E.L., Panchal, S.C., Abraham, V.C., Algire, M.A., Shi, Y., Olson, A.M., et al. (2019). PARP1 Trapping by PARP Inhibitors Drives Cytotoxicity in Both Cancer Cells and Healthy Bone Marrow. Mol Cancer Res 17, 409–419.

Hopkins, T.A., Shi, Y., Rodriguez, L.E., Solomon, L.R., Donawho, C.K., DiGiammarino, E.L., Panchal, S.C., Wilsbacher, J.L., Gao, W., Olson, A.M., et al. (2015). Mechanistic Dissection of PARP1 Trapping and the Impact on In Vivo Tolerability and Efficacy of PARP Inhibitors. Mol Cancer Res 13, 1465–1477.

Huseyin, M.K., and Klose, R.J. (2021). Live-cell single particle tracking of PRC1 reveals a highly dynamic system with low target site occupancy. Nat Commun 12, 887.

Jha, A., and Hansen, A.S. (2022). A Protocol for Studying Transcription FactorTranscription factors Dynamics Using Fast Single-Particle TrackingSingle-particle tracking and Spot-On Model-Based Analysis. In Chromatin: Methods and Protocols, J. Horsfield, and J. Marsman, eds. (New York, NY: Springer US), pp. 151–174.

Johansson, M. (1999). A human poly(ADP-ribose) polymerase gene family (ADPRTL): cDNA cloning of two novel poly(ADP-ribose) polymerase homologues. Genomics 57, 442–445.

Kim, M.Y., Mauro, S., Gevry, N., Lis, J.T., and Kraus, W.L. (2004). NAD+-dependent modulation of chromatin structure and transcription by nucleosome binding properties of PARP-1. Cell 119, 803–814.

Kozlowski, M. (2014). The molecular mechanism of PARP1 activation and its downstream roles in ALC1-regulated transcription. . In Physiologische Chemie (München, Germany., Ludwig Maximilians Universität).

Kraus, W.L. (2008). Transcriptional control by PARP-1: chromatin modulation, enhancer-binding, coregulation, and insulation. Curr Opin Cell Biol 20, 294–302.

Krishnakumar, R., Gamble, M.J., Frizzell, K.M., Berrocal, J.G., Kininis, M., and Kraus, W.L. (2008). Reciprocal binding of PARP-1 and histone H1 at promoters specifies transcriptional outcomes. Science 319, 819–821.

Krishnakumar, R., and Kraus, W.L. (2010). PARP-1 regulates chromatin structure and transcription through a KDM5B-dependent pathway. Mol Cell 39, 736–749.

LaFargue, C.J., Dal Molin, G.Z., Sood, A.K., and Coleman, R.L. (2019). Exploring and comparing adverse events between PARP inhibitors. Lancet Oncol 20, e15–e28.

Langelier, M.F., Planck, J.L., Roy, S., and Pascal, J.M. (2012). Structural basis for DNA damage-dependent poly(ADP-ribosyl)ation by human PARP-1. Science 336, 728–732.

Langelier, M.F., Servent, K.M., Rogers, E.E., and Pascal, J.M. (2008). A third zinc-binding domain of human poly(ADP-ribose) polymerase-1 coordinates DNA-dependent enzyme activation. J Biol Chem 283, 4105–4114.

Le, Y., Miller, J.L., and Sauer, B. (1999). GFPcre fusion vectors with enhanced expression. Anal Biochem 270, 334–336.

Lever, M.A., Th’ng, J.P., Sun, X., and Hendzel, M.J. (2000). Rapid exchange of histone H1.1 on chromatin in living human cells. Nature 408, 873–876.

Liu, L., Kong, M., Gassman, N.R., Freudenthal, B.D., Prasad, R., Zhen, S., Watkins, S.C., Wilson, S.H., and Van Houten, B. (2017). PARP1 changes from three-dimensional DNA damage searching to one-dimensional diffusion after auto-PARylation or in the presence of APE1. Nucleic Acids Res 45, 12834–12847.

Lord, C.J., and Ashworth, A. (2012). The DNA damage response and cancer therapy. Nature 481, 287–294.

Los, G.V., Encell, L.P., McDougall, M.G., Hartzell, D.D., Karassina, N., Zimprich, C., Wood, M.G., Learish, R., Ohana, R.F., Urh, M., et al. (2008). HaloTag: a novel protein labeling technology for cell imaging and protein analysis. ACS Chem Biol 3, 373–382.

Lupey-Green, L.N., Caruso, L.B., Madzo, J., Martin, K.A., Tan, Y., Hulse, M., and Tempera, I. (2018). PARP1 Stabilizes CTCF Binding and Chromatin Structure To Maintain Epstein-Barr Virus Latency Type. J Virol 92.

Mahadevan, J., Bowerman, S., and Luger, K. (2019a). Quantitating repair protein accumulation at DNA lesions: Past, present, and future. DNA Repair (Amst) 81, 102650.

Mahadevan, J., Rudolph, J., Jha, A., Tay, J.W., Dragavon, J., Grumstrup, E.M., and Luger, K. (2019b). Q-FADD: A Mechanistic Approach for Modeling the Accumulation of Proteins at Sites of DNA Damage. Biophys J 116, 2224–2233.

Menissier de Murcia, J., Ricoul, M., Tartier, L., Niedergang, C., Huber, A., Dantzer, F., Schreiber, V., Ame, J.C., Dierich, A., LeMeur, M., et al. (2003). Functional interaction between PARP-1 and PARP-2 in chromosome stability and embryonic development in mouse. EMBO J 22, 2255–2263.

Messner, S., Altmeyer, M., Zhao, H., Pozivil, A., Roschitzki, B., Gehrig, P., Rutishauser, D., Huang, D., Caflisch, A., and Hottiger, M.O. (2010). PARP1 ADP-ribosylates lysine residues of the core histone tails. Nucleic Acids Res 38, 6350–6362.

Michelena, J., Lezaja, A., Teloni, F., Schmid, T., Imhof, R., and Altmeyer, M. (2018). Analysis of PARP inhibitor toxicity by multidimensional fluorescence microscopy reveals mechanisms of sensitivity and resistance. Nat Commun 9, 2678.

Misteli, T., Gunjan, A., Hock, R., Bustin, M., and Brown, D.T. (2000). Dynamic binding of histone H1 to chromatin in living cells. Nature 408, 877–881.

Mortusewicz, O., Ame, J.C., Schreiber, V., and Leonhardt, H. (2007). Feedback-regulated poly(ADP-ribosyl)ation by PARP-1 is required for rapid response to DNA damage in living cells. Nucleic Acids Res 35, 7665–7675.

Murai, J., Huang, S.Y., Das, B.B., Renaud, A., Zhang, Y., Doroshow, J.H., Ji, J., Takeda, S., and Pommier, Y. (2012). Trapping of PARP1 and PARP2 by Clinical PARP Inhibitors. Cancer Res 72, 5588–5599.

Murai, J., Huang, S.Y., Renaud, A., Zhang, Y., Ji, J., Takeda, S., Morris, J., Teicher, B., Doroshow, J.H., and Pommier, Y. (2014). Stereospecific PARP trapping by BMN 673 and comparison with olaparib and rucaparib. Mol Cancer Ther 13, 433–443.

Muthurajan, U.M., Hepler, M.R., Hieb, A.R., Clark, N.J., Kramer, M., Yao, T., and Luger, K. (2014). Automodification switches PARP-1 function from chromatin architectural protein to histone chaperone. Proc Natl Acad Sci U S A 111, 12752–12757.

Nalabothula, N., Al-jumaily, T., Eteleeb, A.M., Flight, R.M., Xiaorong, S., Moseley, H., Rouchka, E.C., and Fondufe-Mittendorf, Y.N. (2015). Genome-Wide Profiling of PARP1 Reveals an Interplay with Gene Regulatory Regions and DNA Methylation. PLoS One 10, e0135410.

Noordermeer, S.M., and van Attikum, H. (2019). PARP Inhibitor Resistance: A Tug-of-War in BRCA-Mutated Cells. Trends Cell Biol 29, 820–834.

Poirier, G.G., de Murcia, G., Jongstra-Bilen, J., Niedergang, C., and Mandel, P. (1982). Poly(ADP-ribosyl)ation of polynucleosomes causes relaxation of chromatin structure. Proc Natl Acad Sci U S A 79, 3423–3427.

Pommier, Y., O’Connor, M.J., and de Bono, J. (2016). Laying a trap to kill cancer cells: PARP inhibitors and their mechanisms of action. Sci Transl Med 8, 362ps317.

Ray Chaudhuri, A., and Nussenzweig, A. (2017). The multifaceted roles of PARP1 in DNA repair and chromatin remodelling. Nat Rev Mol Cell Biol 18, 610–621.

Ronson, G.E., Piberger, A.L., Higgs, M.R., Olsen, A.L., Stewart, G.S., McHugh, P.J., Petermann, E., and Lakin, N.D. (2018). PARP1 and PARP2 stabilise replication forks at base excision repair intermediates through Fbh1-dependent Rad51 regulation. Nat Commun 9, 746.

Rose, M., Burgess, J.T., O’Byrne, K., Richard, D.J., and Bolderson, E. (2020). PARP Inhibitors: Clinical Relevance, Mechanisms of Action and Tumor Resistance. Front Cell Dev Biol 8, 564601.

Rudolph, J., Jung, K., and Luger, K. (2022). Inhibitors of PARP: Number crunching and structure gazing. Proc Natl Acad Sci U S A 119, e2121979119.

Rudolph, J., Mahadevan, J., Dyer, P., and Luger, K. (2018). Poly(ADP-ribose) polymerase 1 searches DNA via a ‘monkey bar’ mechanism. Elife 7.

Rudolph, J., Mahadevan, J., and Luger, K. (2020). Probing the Conformational Changes Associated with DNA Binding to PARP1. Biochemistry 59, 2003–2011.

Rudolph, J., Muthurajan, U.M., Palacio, M., Mahadevan, J., Roberts, G., Erbse, A.H., Dyer, P.N., and Luger, K. (2021a). The BRCT domain of PARP1 binds intact DNA and mediates intrastrand transfer. Mol Cell 81, 4994–5006 e4995.

Rudolph, J., Roberts, G., and Luger, K. (2021b). Histone Parylation factor 1 contributes to the inhibition of PARP1 by cancer drugs. Nat Commun 12, 736.

Saha, A., and Dalal, Y. (2021). A glitch in the snitch: the role of linker histone H1 in shaping the epigenome in normal and diseased cells. Open Biol 11, 210124.

Schmidt, J.C., Zaug, A.J., and Cech, T.R. (2016). Live Cell Imaging Reveals the Dynamics of Telomerase Recruitment to Telomeres. Cell 166, 1188–1197 e1189.

Serge, A., Bertaux, N., Rigneault, H., and Marguet, D. (2008). Dynamic multiple-target tracing to probe spatiotemporal cartography of cell membranes. Nat Methods 5, 687–694.

Shao, Z., Lee, B.J., Rouleau-Turcotte, E., Langelier, M.F., Lin, X., Estes, V.M., Pascal, J.M., and Zha, S. (2020). Clinical PARP inhibitors do not abrogate PARP1 exchange at DNA damage sites in vivo. Nucleic Acids Res 48, 9694–9709.

Sprague, B.L., Pego, R.L., Stavreva, D.A., and McNally, J.G. (2004). Analysis of binding reactions by fluorescence recovery after photobleaching. Biophys J 86, 3473–3495.

Strickfaden, H., McDonald, D., Kruhlak, M.J., Haince, J.F., Th’ng, J.P.H., Rouleau, M., Ishibashi, T., Corry, G.N., Ausio, J., Underhill, D.A., et al. (2016). Poly(ADP-ribosyl)ation-dependent Transient Chromatin Decondensation and Histone Displacement following Laser Microirradiation. J Biol Chem 291, 1789–1802.

Sukhanova, M.V., Abrakhi, S., Joshi, V., Pastre, D., Kutuzov, M.M., Anarbaev, R.O., Curmi, P.A., Hamon, L., and Lavrik, O.I. (2016). Single molecule detection of PARP1 and PARP2 interaction with DNA strand breaks and their poly(ADP-ribosyl)ation using high-resolution AFM imaging. Nucleic Acids Res 44, e60.

Thorsell, A.G., Ekblad, T., Karlberg, T., Low, M., Pinto, A.F., Tresaugues, L., Moche, M., Cohen, M.S., and Schuler, H. (2017). Structural Basis for Potency and Promiscuity in Poly(ADP-ribose) Polymerase (PARP) and Tankyrase Inhibitors. J Med Chem 60, 1262–1271.

Tokunaga, M., Imamoto, N., and Sakata-Sogawa, K. (2008). Highly inclined thin illumination enables clear single-molecule imaging in cells. Nat Methods 5, 159–161.

Watanabe, N., and Mitchison, T.J. (2002). Single-molecule speckle analysis of actin filament turnover in lamellipodia. Science 295, 1083–1086.

Xi, L., Schmidt, J.C., Zaug, A.J., Ascarrunz, D.R., and Cech, T.R. (2015). A novel two-step genome editing strategy with CRISPR-Cas9 provides new insights into telomerase action and TERT gene expression. Genome Biol 16, 231.

Yi, M., Dong, B., Qin, S., Chu, Q., Wu, K., and Luo, S. (2019). Advances and perspectives of PARP inhibitors. Exp Hematol Oncol 8, 29.

Youmans, D.T., Schmidt, J.C., and Cech, T.R. (2018). Live-cell imaging reveals the dynamics of PRC2 and recruitment to chromatin by SUZ12-associated subunits. Genes Dev 32, 794–805.

Zandarashvili, L., Langelier, M.F., Velagapudi, U.K., Hancock, M.A., Steffen, J.D., Billur, R., Hannan, Z.M., Wicks, A.J., Krastev, D.B., Pettitt, S.J., et al. (2020). Structural basis for allosteric PARP-1 retention on DNA breaks. Science 368.

Zentout, S., Smith, R., Jacquier, M., and Huet, S. (2021). New Methodologies to Study DNA Repair Processes in Space and Time Within Living Cells. Front Cell Dev Biol 9, 730998.

