## Supplementary figs and legends for "Dynamics of endogenous PARP1 and PARP2 during DNA damage revealed by live-cell single-molecule imaging"

### Supp figure 1

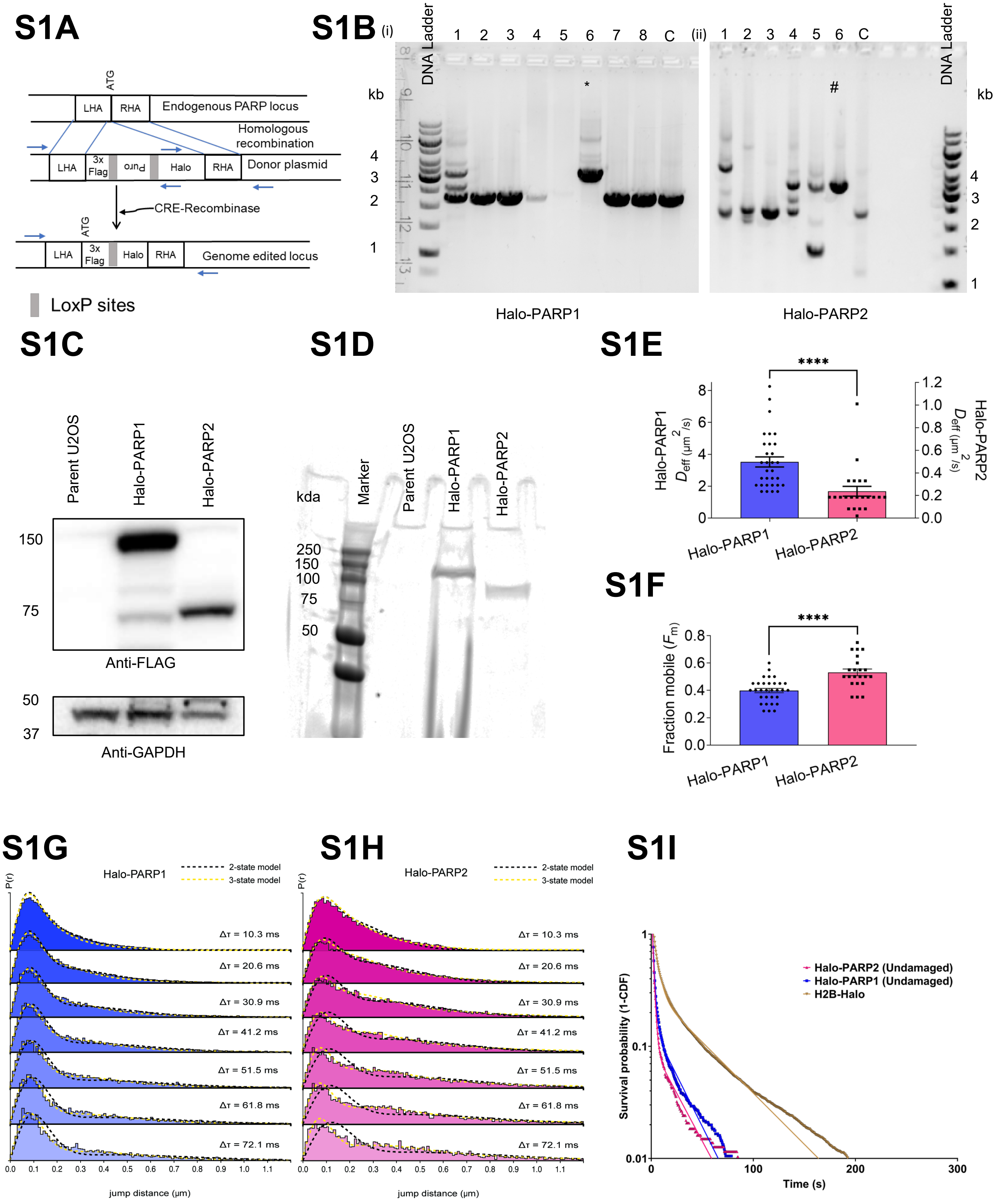

#### Supp figure 2

### S2A

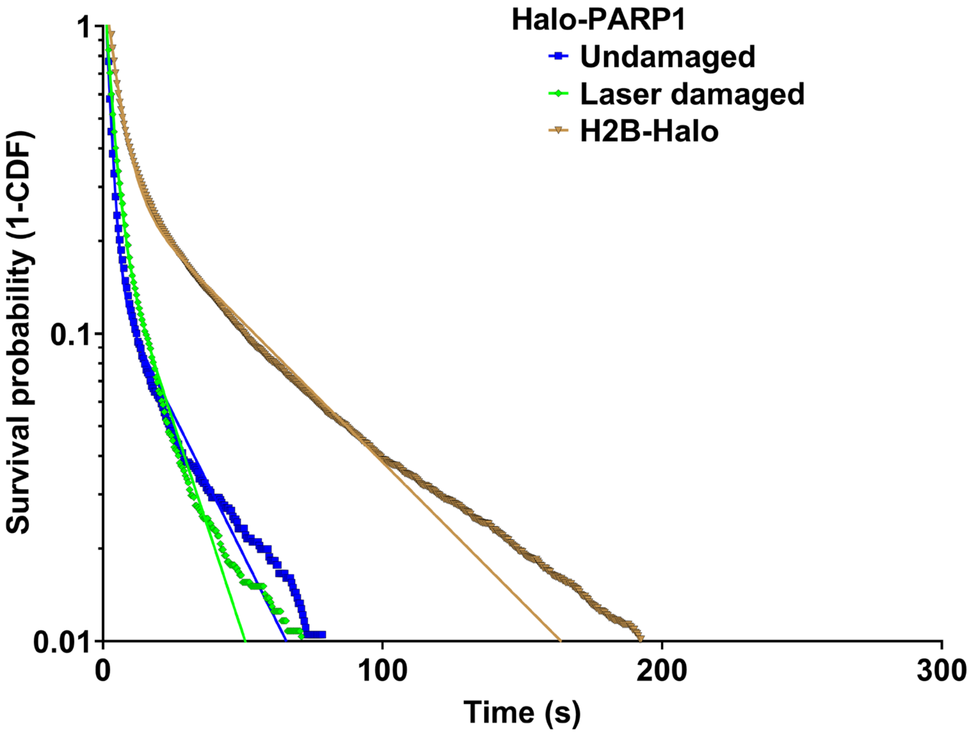


### S2B

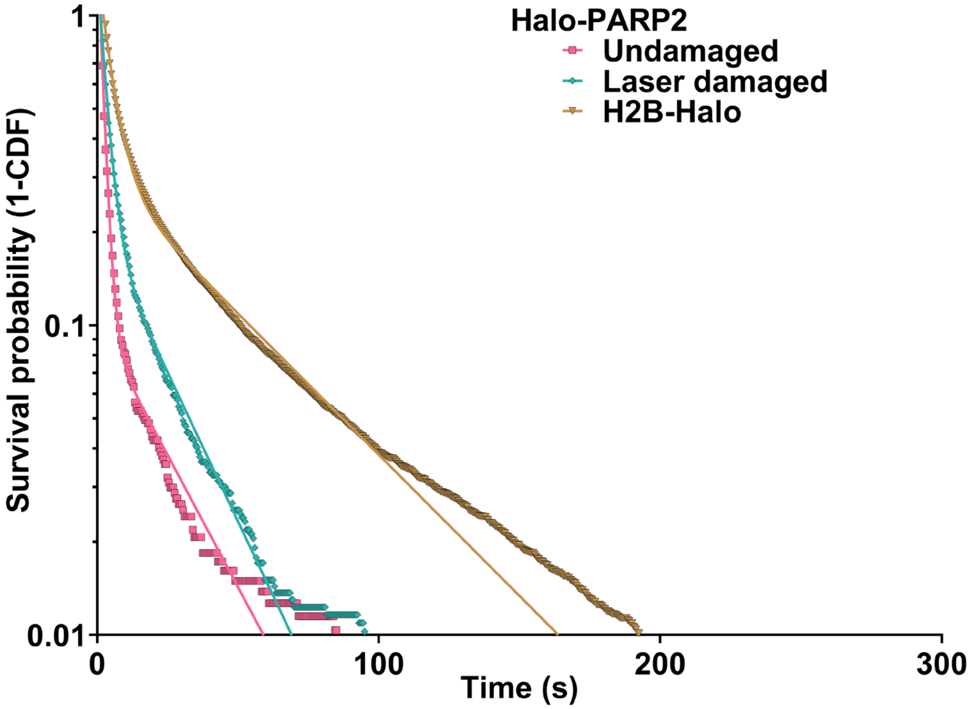

### Supp figure 3

## S3A

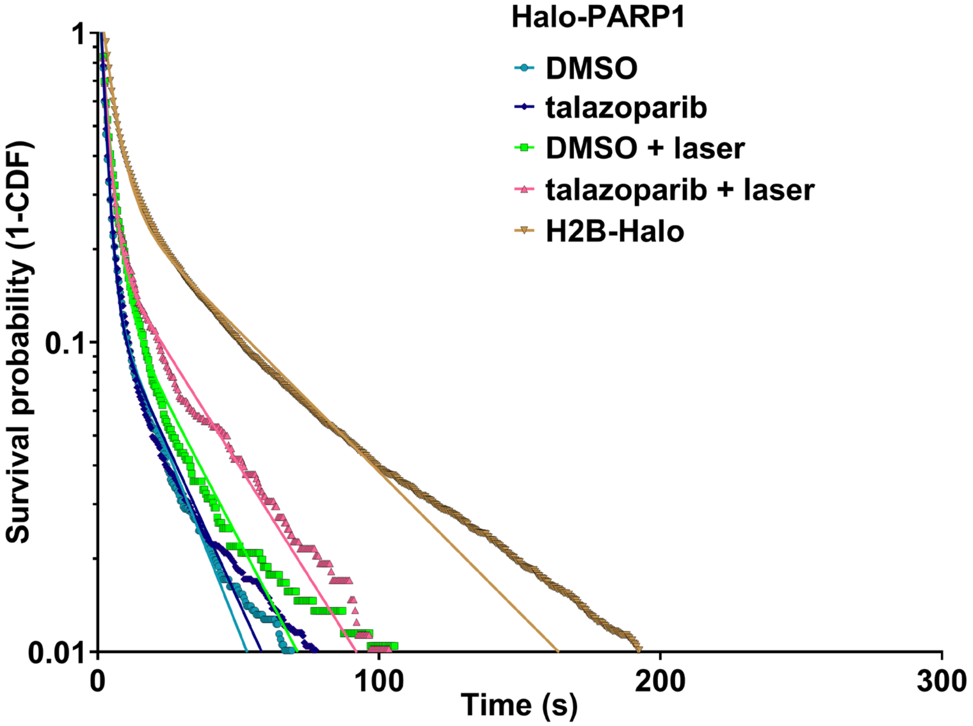


## S3B

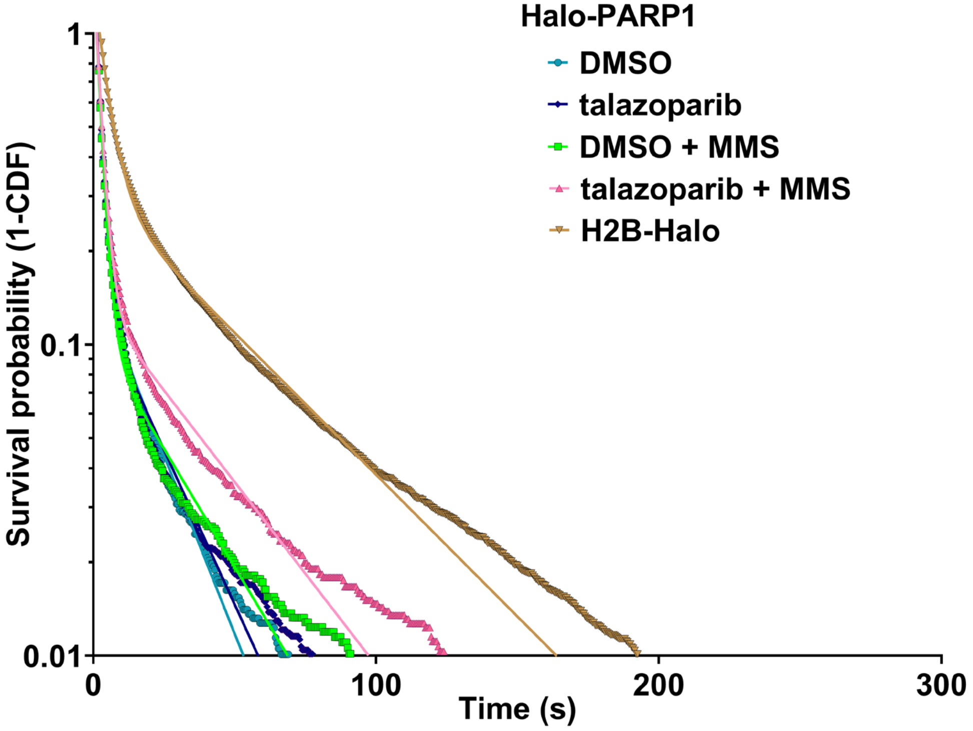

### Supp figure 4

## S4A

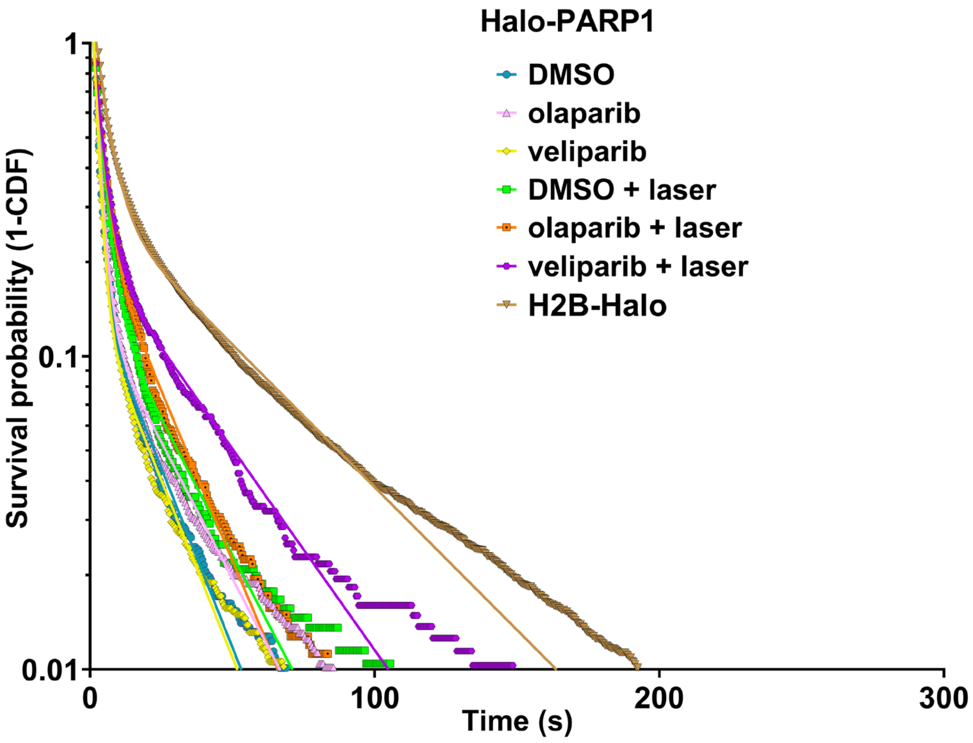


## S4B

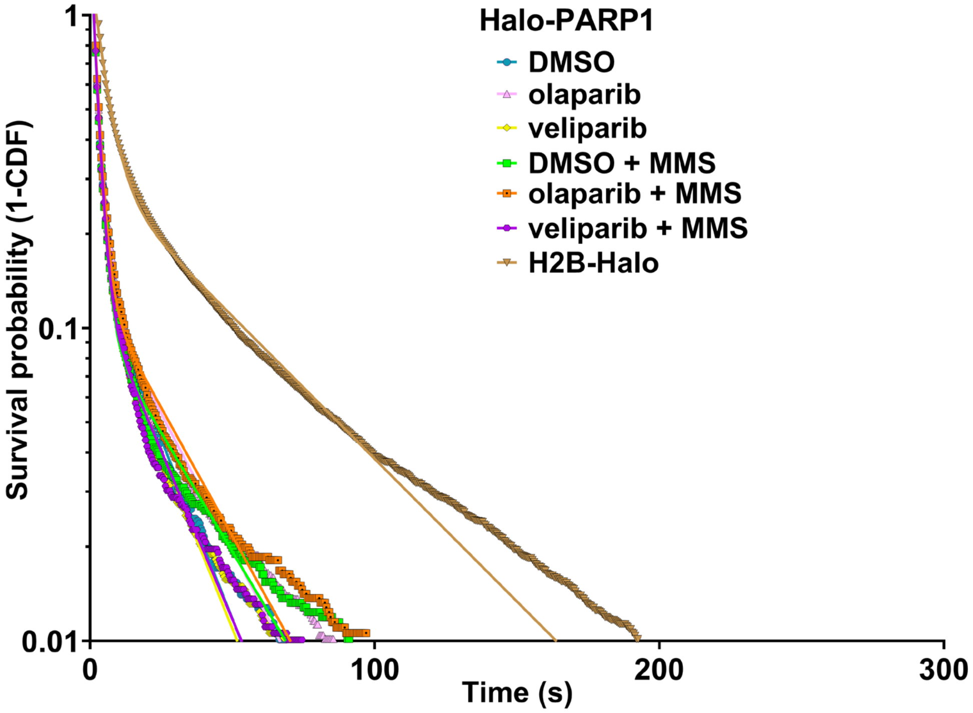

### Supp figure 5

## S5A

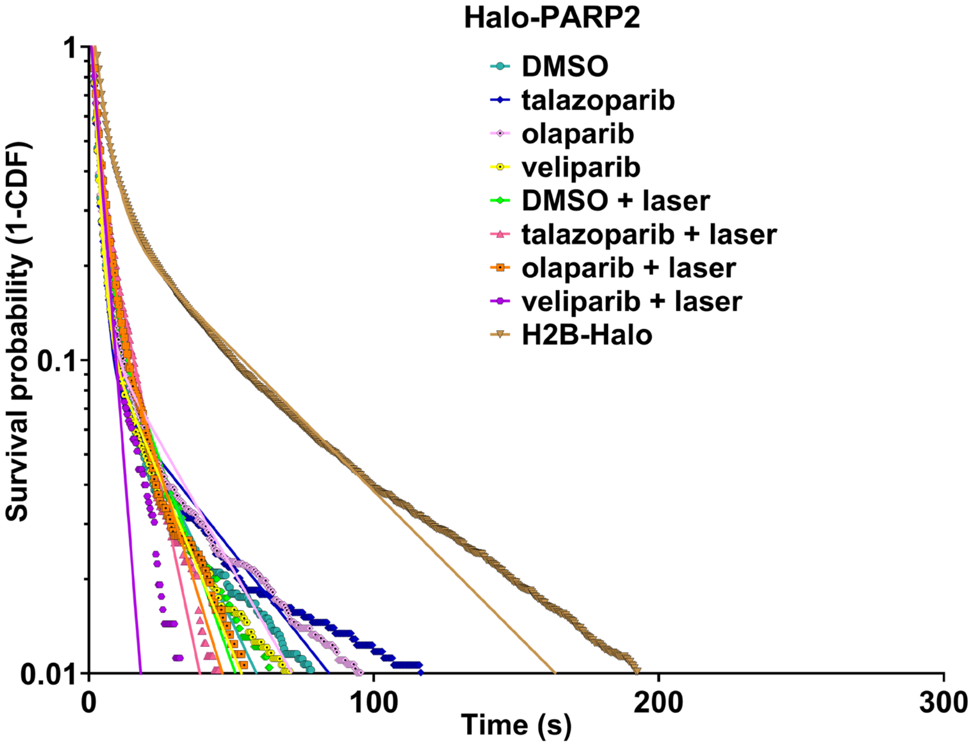


## S5B

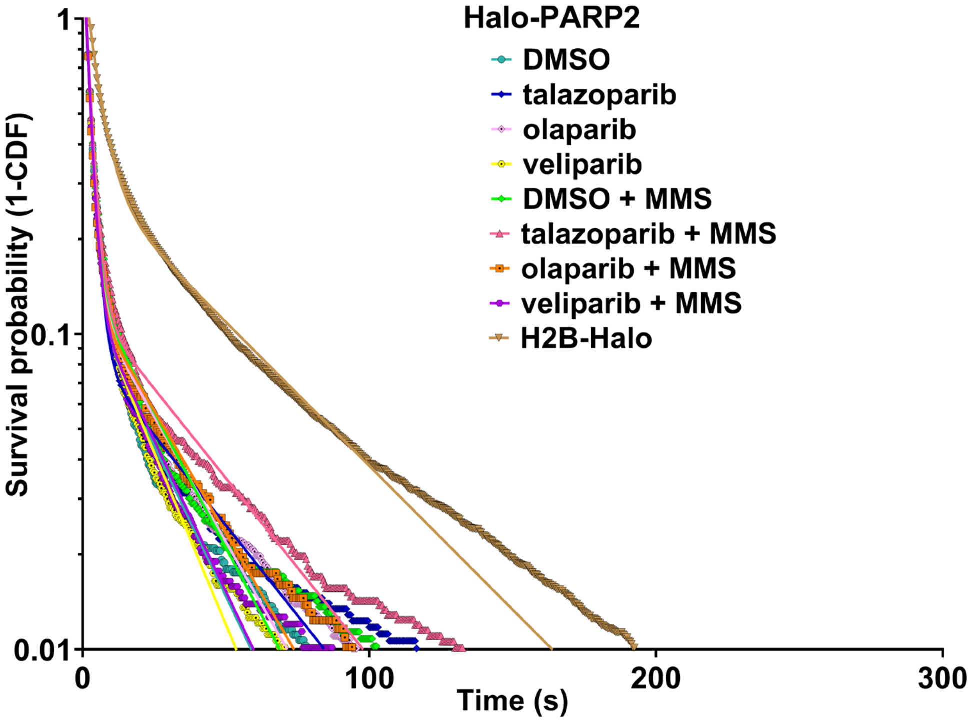

**Figure S1. Live-cell single molecule microscopy reveals fraction of stably bound PARP1 and PARP2 in undamaged cells (Related to Figure 1)**

- **A.** CRISPR/Cas9 based genome editing scheme for homozygously inserting 3X Flag HaloTag into the N-termini of PARP1 and PARP2 genes. A donor plasmid consisting of the 3X Flag HaloTag, puromycin resistance cassette and left and right homology arm regions was introduced to enable homologous recombination (HR). Further, EGFP-Cre recombinase was ectopically expressed to excise the puromycin cassette in the population of puromycin selected cells that have undergone HR. This population of EGFP+ cells was used to establish single cell clones and thereby genome edited Halo-PARP1 and Halo-PARP2 U2OS cell lines.
- **B.** Agarose gels loaded with PCR products amplified using genomic DNA from single cell clones generated after genome editing as template and primers designed outside the left and right homology arm (LHA and RHA) as indicated in Figure S1A. Appropriate genome editing yielded product sizes of 3.2 kb for Halo-PARP1 (lane 6 in i, indicated by *) and 3.4 kb for Halo-PARP2 (lane 6 in ii indicated by #). C refers to control lanes loaded with PCR amplified products from unedited parent U2OS cells using the same primers.
- **C.** Immunoblots using anti-Flag antibody indicating protein levels of Flag-Halo-PARP1/2 in parent and genome edited cell lines. GAPDH was used as a loading control.
- **D.** SDS-PAGE gels imaged using the 647 nm channel, showing protein levels of Halo-PARP1 and Halo-PARP2. Halo tagged proteins in whole cell extracts of genome edited U2OS cells were labeled with the HaloTag ligand, JF646.
- **E. F.** Bar graphs depicting the mean ± standard error (SEM) of effective diffusion coefficient ($D_{\text{eff}}$ ) (in E) and fraction mobile ($F_{\text{m}}$) (in F) of Halo-PARP1 and Halo-PARP2 for ≥ 22 cells from ≥ 3 independent experiments, inferred from Q-FADD analysis performed on individual cells subjected to bulk laser microirradiation (represented by dots). Statistical differences between the two groups were determined using unpaired t-test.
- **G. H.** Two-state (black dashed line) and three-state (yellow dashed line) model fits overlaid on pooled jump length (μm) histograms for multiple time delays ($\Delta\tau$) for Halo-PARP1 (in G) and Halo-PARP2 (in H). Data from all cells in one representative replicate dataset are included in each histogram.
- **I.** A semi-log plot of data presented in Figure 1F.

**Figure S2. Majority of PARP1 and PARP2 molecules diffuse freely at laser-induced DNA lesions (Related to Figure 2)**

- **A. B.** Semi-log plots of data presented in Figure 2D (in A) and 2E (in B)

**Figure S3. An efficient PARP trapping agent, talazoparib, increases the retention time of only a small fraction of stably binding PARP1 molecules at damage sites (Related to Figure 3)**

- **A. B.** Semi-log plots of data presented in Figure 3D (in A) and 3E (in B)

**Figure S4. Weaker PARP trapping agents, olaparib and veliparib, exert distinct effects on the retention time of stably binding PARP1 molecules (Related to Figure 4)**

- **A. B.** Semi-log plots of data presented in Figure 4D (in A) and 4E (in B)

**Figure S5. Trapping of stably binding PARP2 molecules mediated by talazoparib does not require induction of DNA damage (Related to Figure 5)**

**A. B.** Semi-log plots of data presented in Figure 5B (in A) and 5C (in B)
